# HDA6-mediated histone deacetylation restricts premature tapetal PCD to maintain male fertility during early flowering in Arabidopsis

**DOI:** 10.64898/2026.09.17.752293

**Authors:** Kangwei Hu, Xun Weng, Ling Zhou, Junyu Chen, Cheng-Guo Duan, Lifeng Zhao, He Yan, Hao Wang

**Affiliations:** Department of Cell and Developmental Biology, College of Life Sciences, Guangdong Provincial Key Laboratory for the Developmental Biology and Environmental Adaption of Agricultural Organisms, South China Agricultural University, Guangzhou 510642, China; State Key Laboratory of Plant Trait Design, CAS Center for Excellence in Molecular Plant Sciences, Chinese Academy of Sciences, Shanghai 200032, China; School of Agriculture and Biotechnology, Shenzhen Campus of Sun Yat-sen University, Sun Yat-sen University, Shenzhen 518107, China

**Keywords:** Histone deacetylation, tapetal programmed cell death, male gametophyte development, male fertility, *Arabidopsis thaliana*

## Abstract

Histone deacetylation regulates plant development, yet its role in male gametophyte development and fertility remains unclear. Here, we show that histone deacetylase 6 (HDA6), an RPD3-like histone deacetylase, epigenetically regulates timely tapetal programmed cell death (PCD) and male fertility, predominantly during early flowering in Arabidopsis. Loss of HDA6 in *axe1-4* causes early-stage male sterility followed by fertility recovery, accompanied by premature tapetal PCD, impaired pollen development and defective pollen wall formation. HDA6 interacts with squamosa promoter binding protein-like 8 (SPL8) to repress *Arabidopsis NAC domain containing protein 087* (*ANAC087*) through H3K9/K14 deacetylation. In *axe1-4*, *ANAC087* derepression activates *cysteine endopeptidase 1* (*CEP1*), triggering precocious tapetal PCD. Genetic suppression of *ANAC087* or *CEP1* restores male fertility. Thus, HDA6-SPL8 establishes a stage-dependent epigenetic checkpoint that restrains the *ANAC087*-*CEP1* cell death pathway and ensures timely tapetal PCD. Our study reveals how chromatin regulation integrates with developmental timing to safeguard plant male fertility.

## Introduction

Plant male fertility is a complex developmental trait essential for sexual reproduction and seed formation, and its disruption can lead to severe substantial yield loss (*1, 2*). In flowering plants, male gametophyte development and fertility depend on the precise differentiation and timely degeneration of the tapetum, the innermost somatic cell layer of the anther (*3, 4*). The tapetum provides nutrients, enzymes and structural molecules for microspore development and pollen wall formation. Its programmed cell death (PCD) must be spatiotemporally tightly controlled, as either premature or delayed tapetal PCD disrupts pollen maturation and ultimately causes male sterility (*5, 6*). Although a number of transcription factors and proteases have been implicated in regulating tapetal PCD, the epigenetic regulatory mechanisms that govern the developmental timing of this process remain largely unexplored (*7–9*).

Histone deacetylases (HDAs) are key epigenetic regulators that remove acetyl groups from histone tails, thereby promoting chromatin condensation and transcriptional repression (*10, 11*). In Arabidopsis, HDA6, a class I RPD3-like deacetylase, has been implicated in a wide range of developmental and physiological processes including the regulation of flowering time, leaf senescence and versatile stress responses (*12–16*). HDA6 can also be recruited by specific transcription factors to silence target genes, indicating that it could operate in tissue-specific transcriptional regulations (*17*). However, its role in plant male gametophyte development and fertility remains poorly understood.

The regulation of tapetal PCD involves a complex transcriptional network (*18, 19*). Within the network, cysteine protease 1 (CEP1) functions as a key executor of tapetal PCD, as its overexpression triggers premature tapetal degeneration and male sterility (*20*). Upstream of *CEP1*, the Arabidopsis NAC transcription factor (ANAC087) activates *CEP1* expression and has also been implicated in leaf senescence, pointing to a potential role in controlling the onset of tapetal PCD (*21*). However, how this pathway is repressed during early anther development to prevent premature tapetal PCD, and ensure proper male gametophyte development and fertility remains unknown. Elucidating the upstream regulatory mechanisms that constrain ANAC087-*CEP1* signaling may therefore provide critical insight into the transcriptional control of tapetal PCD progression and the maintenance of male fertility. In parallel, increasing evidence indicates that transcription factors can recruit histone-modifying enzymes to specific genomic loci, thereby facilitating precise spatiotemporal control of gene expression (*22, 23*). Given its essential role in anther and pollen development, squamosa promoter binding protein-like 8 (SPL8) is well positioned to mediate such chromatin-associated regulation, consistent with the tapetal defects and male sterility observed in *spl8* mutants (*24, 25*). Nevertheless, whether SPL8 exerts its function through chromatin-based regulation, including potential cooperation with histone modifiers such as HDA6, remains to be determined.

Here, we show that HDA6 interacts with SPL8 to repress *ANAC087* expression by reducing H3K9/K14 acetylation at its promoter. Loss of HDA6 leads to derepression of the *ANAC087*-*CEP1* module, resulting in premature tapetal PCD, defective pollen wall formation and male sterility. Notably, genetic suppression of either *ANAC087* or *CEP1* in hda6 mutants rescues these fertility defects, demonstrating the functional importance of this pathway. Our findings reveal a chromatin-based epigenetic regulatory mechanism that fine-tunes tapetal PCD timing to ensure proper male gametophyte development and fertility, with potential implications for crop breeding.

## Results

### HDA6 is required for early male fertility and proper pollen coat formation in Arabidopsis

The Arabidopsis *axe1-4* mutant, an EMS-induced *hda6* allele carrying a mutation within the deacetylase domain, exhibited a marked reduction in seed set in the basal siliques formed during the early reproductive stage and delayed flowering (Fig. 1A and Fig. S1A). To test whether this fertility defect was attributable to the loss of HDA6 function, we expressed *pHDA6::HDA6-myc* and *pHDA6::HDA6-GFP* in the *axe1-4* background. Both transgenes fully complemented the early fertility defect of *axe1-4* (Fig. 1A and Fig. S1). We next introduced targeted mutations at sites adjacent to the *axe1-4* mutation site in *HDA6* in the wild-type (WT) Arabidopsis (Col-0) by CRISPR/Cas9, thereby generating the independent alleles *hda6-8* and *hda6-43* mutants with the aim of recapitulating the *axe1-4* defects (Fig. S2A). Both mutants showed early fertility defects and delayed flowering comparable to those of *axe1-4* (Fig. 1A and Fig. S2B-D). These results demonstrate that the early fertility defect observed in *axe1-4* is caused by loss of HDA6 function.

**Figure 1.**
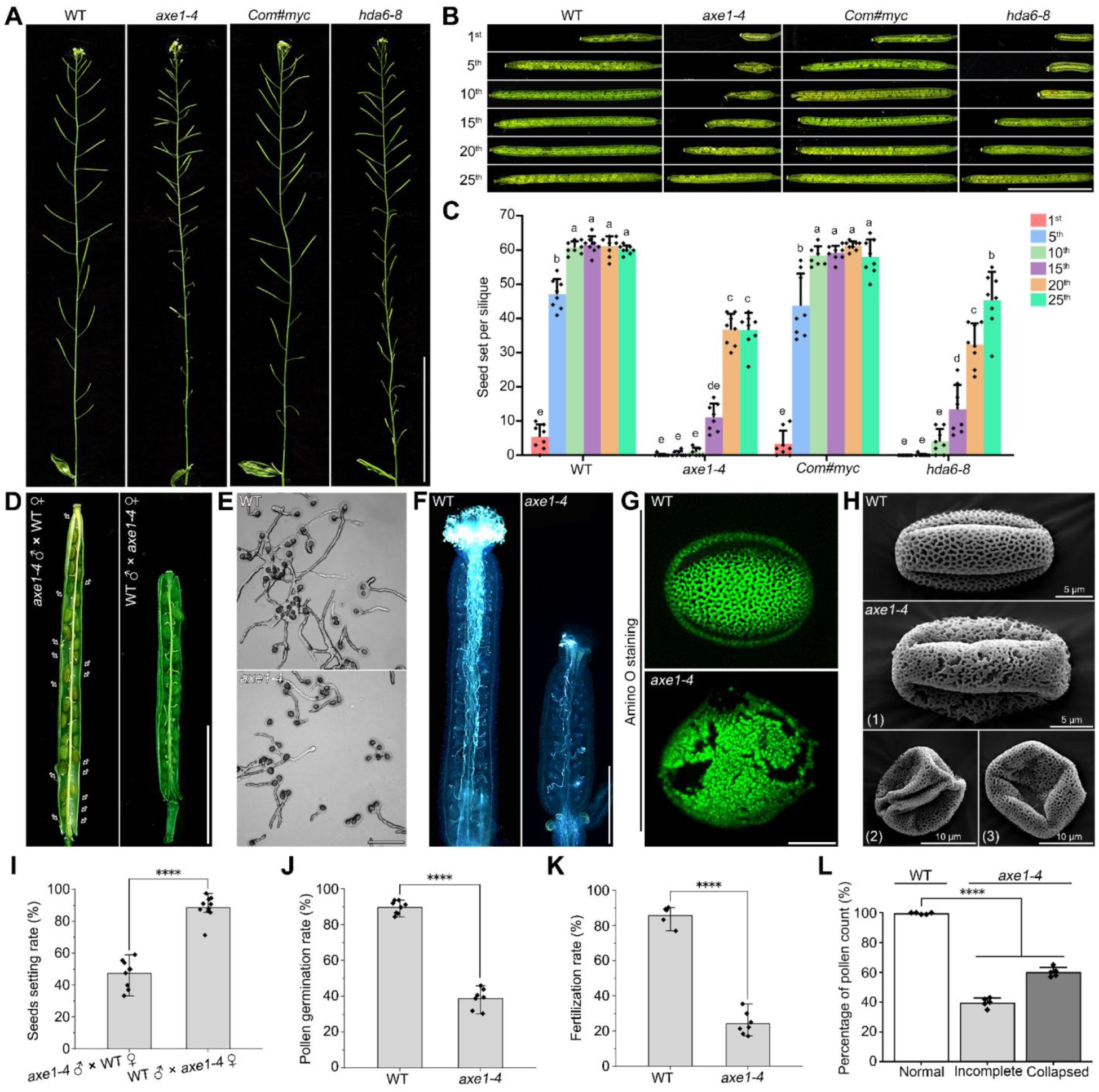
Loss of HDA6 function impairs pollen fertility and male gametophyte development in Arabidopsis. (**A**) Representative images of primary inflorescences, excised below the last cauline leaf of Arabidopsis wild type (Col-0), *axe1-4*, the complemented line *Com#myc* and *hda6-8.* (**B**) Representative images of siliques corresponding to the 1^st^, 5^th^, 10^th^, 15^th^, 20^th^ and 25^th^ flowers formed on the primary inflorescences of the indicated genotypes. (**C**) Statistical quantification of seed number per silique at the indicated flower positions on the primary inflorescences of the indicated genotypes (n = 8). Different letters indicate significant differences among groups as determined by one-way ANOVA followed by Tukey’s multiple comparison test (*P* < 0.05). (**D**) Seeds setting after cross-pollination between WT and *axe1-4*. (**E**) Representative images of *in vitro* pollen germination in WT and *axe1-4.* (**F**) Representative images of *in vivo* pollen germination and pollen tube growth in WT and *axe1-4.* (**G**) Amino O staining of mature pollen grains from WT and *axe1-4*. (**H**) Scanning electron microscopy imaging of mature pollen of Col-0 and *axe1-4.* Representative images of *axe1-4* pollen grains exhibiting incomplete exine formation (1) and collapsed morphology (2 and 3) are shown. (**I**) Calculation of seed-setting rate after reciprocal crosses between WT and *axe1-4* (n ≥ 8). (**J**) Quantification of *in vitro* pollen germination rate in WT and *axe1-4* (n ≥ 300). (**K**) Quantification of successful *in vivo* fertilization rate in WT and *axe1-4* (n ≥ 6). (**L**) Quantification of pollen grain morphology in WT and *axe1-4*, classified as normal, incomplete or collapsed (n ≥ 100). All of the samples in (**D**-**L**) were collected from early flowers before the 15^th^ flower. Values are means ±SD. Asterisks indicate statistically significant differences: \**P* < 0.05, \*\**P* < 0.01, \*\*\**P* < 0.001 and \*\*\*\**P* < 0.0001. Scale bars = 8 cm in (**A**), 10 mm in (**B**), 4 mm in (**D**), 150 μm in (**E**), 1 mm in (**F**) and 10 μm in (**G**).

To further define the stage specificity of the fertility defect in *axe1-4*, we next examined seed set in successive siliques along the primary inflorescence. Siliques derived from the 1^st^, 5^th^, 10^th^, 15^th^, 20^th^ and 25^th^ flowers to open were sequentially marked, and seed number was subsequently quantified. In the WT, seed set increased rapidly from approximately five seeds in the 1^st^ silique to approximately 48 seeds in the 5^th^, and then plateaued at around 60 seeds per silique from the 10^th^ through the 25^th^ siliques (Fig. 1B and C), consistent with previous studies (*26, 27*). By contrast, *axe1-4* mutant produced almost no seeds in siliques formed up to the 10^th^ flower, indicating a severe fertility defect during the early flowering stage. Although seed set began to recover in the 15^th^ silique and was substantially restored in the 20^th^ and later siliques, the number of seeds per silique remained lower than in WT (Fig. 1B and C). The early reproductive defect was fully rescued by *pHDA6::HDA6-myc*, whereas *hda6-8* largely recapitulated the *axe1-4* phenotype (Fig. 1B and C). These findings indicate that HDA6 is specifically required to maintain fertility during early flowering stage of Arabidopsis.

To better determine whether the fertility defect in *axe1-4* arises from impaired male or/and female gametophytic function, we firstly performed cross pollination between the WT and *axe1-4* at early flowering stage. WT pollen fully restored seed set when applied to *axe1-4* pistils, whereas *axe1-4* pollen reduced seed set to approximately 40% when applied to WT pistils (Fig. 1D and I). Notably, this residual reduction is attributable to decreased ovule number rather than impaired ovule fertility, as mutant ovules remained fully fertile. Loss of *HDA6* was associated with reduced ovule number and shorter pistils without affecting female gametophyte fertility, although the mechanism through which HDA6 regulates female reproductive output remains unclear (Fig. S3A and B). To further determine how the fertility defect in *axe1-4* arises from impaired male gametophytic function, we then assessed the germination ability of mature pollen collected from flowers before the 20^th^ position on the primary inflorescence. *In vitro* pollen germination assays showed that the germination rate of *axe1-4* pollens was approximately 35%, markedly lower than the approximately 90% observed in the WT (Fig. 1E and J). Consistent with this reduction, *in vivo* pollination assays showed that only a subset of *axe1-4* pollen grains germinated normally and achieved successful fertilization (Fig. 1 *F* and *K*). However, the sperm cell formation appeared normal in *axe1-4* compared with WT (Fig. S3C). Thus, the early fertility defect of *axe1-4* is primarily male-derived rather than caused by female gametophytic failure.

In light of the predominantly male gametophytic defect observed in *axe1-4*, we next examined pollen morphology and pollen coat structure in the mutant. Confocal microscopy imaging of Auramine O-stained pollen grains revealed pronounced exine structural disruption in *axe1-4* relative to the WT (Fig. 1G). Consistent with this observation, scanning electron microscopy (SEM) imaging showed that WT pollen grains displayed a characteristic ellipsoidal morphology and regular reticulate exine pattern, whereas *axe1-4* pollen grains were severely abnormal (Fig. 1H). Approximately 60% of *axe1-4* pollen grains were collapsed, while most of the remaining pollens retained an overall structural integrity but exhibited extensive exine damage (Fig. 1L). These results indicate that HDA6 is required for proper pollen coat formation in Arabidopsis.

Given the marked pollen coat structural defects observed in *axe1-4*, we next analyzed the lipid composition of mature pollen grains from the WT and *axe1-4* by gas chromatography-mass spectrometry (GC-MS) (Fig. S4). Compared with WT, *axe1-4* pollen showed a substantial reduction in major wax constituents including the free fatty acids hexadecanoic acid (16:0), octadecanoic acid (18:0) and the alkane nonacosane (C29:0) (Fig. S4). Less abundant long-chain fatty acids including heptacosanoic acid and octacosanoic acid were also reduced, whereas several minor alkane and sterol components were largely unaffected (Fig. S4). These results suggest that loss of *HDA6* broadly reduces the major wax constituents of mature pollen grains and alters pollen coat lipid composition, which may contribute to the defective pollen coat and reduced fertility of *axe1-4*.

### Loss of HDA6 disrupts tapetal PCD timing and male gametophyte maturation

RNA *in situ* hybridization showed that *HDA6* is preferentially expressed in the tapetum of WT anthers, with pronounced accumulation during early stages of anther development (Fig. 2A). To determine the developmental basis of the fertility defects in *axe1-4*, we next examined transverse sections of WT and *axe1-4* anther at successive stages of development. At stage 8, WT anthers contained compact and morphologically intact tapetal cells, and the released microspores were uniformly stained and structurally regular (Fig. 2B). In contrast, stage 8 *axe1-4* anthers exhibited loosely organized, vacuolated tapetal cells, consistent with premature onset of PCD, and some microspores had already become morphologically irregular (Fig. 2C). These abnormalities became more pronounced at stage 9, when *axe1-4* microspores displayed marked vacuolization, severe morphological distortion and apparent defects in sporopollenin deposition (Fig. 2C). By stage 10, while the WT tapetum had only begun normal degeneration, the *axe1-4* tapetum was already highly vacuolated and largely degraded (Fig. 2C). By stage 11, *axe1-4* anthers contained polymorphic pollen grains with extensive vacuolization and disrupted intracellular organization (Fig. 2C).

**Figure 2.**
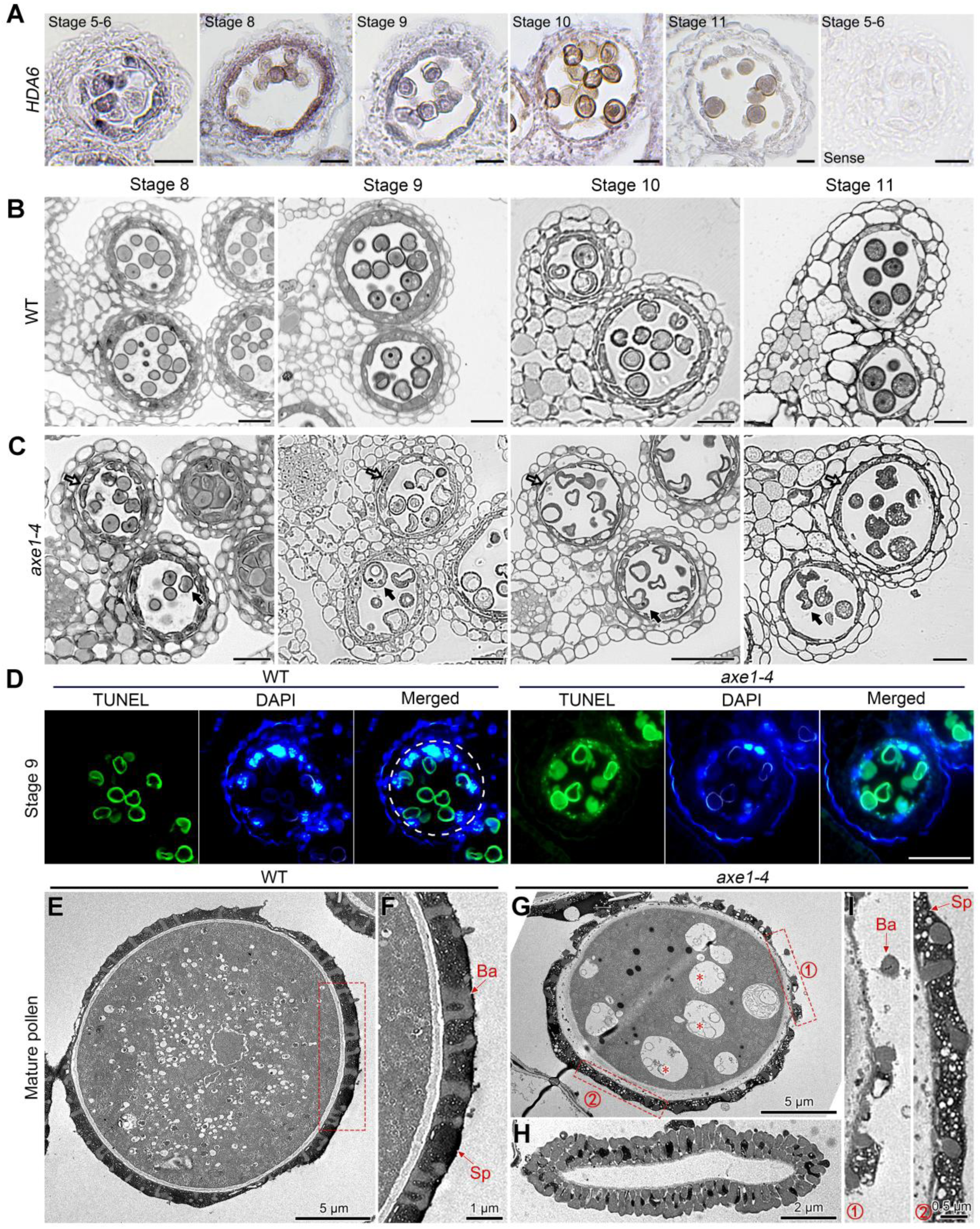
HDA6 is required for proper tapetal PCD and pollen development. (**A**) Representative images of *HDA6* mRNA signaling in WT by *in situ* hybridization assays (n = 6). (**B** and **C**) Representative images of semithin transverse sections of WT (**B**) and *axe1-4* (**C**) anthers at stages 8-11 (n ≥ 3). Hollow arrows indicate abnormal tapetal degeneration and solid arrows show defective microspore or pollen morphology in *axe1-4*. (**D**) Representative images of TUNEL assays analysis of stage 9 anthers of WT and *axe1-4* mutant. TUNEL signal, green; DAPI, blue. White dashed lines indicate the tapetal cell layer (n ≥ 3). (**E***-***I**) Representative images of TEM imaging of stage mature pollen grains from WT and *axe1-4* (n ≥ 6). (**E**) WT pollen grain. Enlarged view of the boxed region in (**F**). (**G**) Representative *axe1-4* pollen grain showing disrupted vacuole defragmentation and defects in wall and coat formation. Asterisks indicate large vacuoles. (**H**) Collapsed *axe1-4* pollen grain without internal cellular contents. (**I**) Enlarged views of the boxed regions in (**G**) showing defective exine structure and reduced sporopollenin accumulation respectively in *axe1-4*. All of the samples were collected from early flowers before the 15^th^ flower. Ba, bacula; Sp, sporopollenin. Scale bars = 20 μm in (**A**), 20 μm in (**B**) and (**C**), and 50 μm in (**D**).

Given the precocious tapetal degeneration observed in *axe1-4*, we then examined whether loss of *HDA6* alters the timing of tapetal PCD. Terminal deoxynucleotidyl transferase-mediated dUTP nick end labeling (TUNEL) assays revealed no signal in the tapetum of WT anthers at stage 9 (Fig. 2D). In contrast, a strong TUNEL-positive signal was already present in the tapetum of *axe1-4* anthers at the same stage, indicating premature initiation of tapetal PCD in the mutant (Fig. 2D). At stage 10, TUNEL signals appeared in the WT tapetum, whereas the signal in *axe1-4* had already begun to weaken, suggesting that tapetal degeneration was nearing completion in the mutant (*SI Appendix*, Fig. S5A). At stage 11, the WT tapetum showed strong TUNEL positivity, while only weak residual signal remained in *axe1-4* (Fig. S5B). By stage 12, no TUNEL signal was detected in either genotype, consistent with complete tapetal degradation (Fig. S5C). These results indicate that HDA6 is required for proper temporal control of tapetal PCD, and that its loss leads to precocious tapetal degeneration likely underlying the pollen defects and male sterility of *axe1-4*.

Consistent with these developmental abnormalities, transmission electron microscopy further revealed severe ultrastructural defects in mature *axe1-4* pollen grains. WT pollen grains were fully developed and exhibited intact pollen walls, uniform sporopollenin deposition and fulfilled with intracellular contents (Fig. 2E and F). By contrast, *axe1-4*pollen grains exhibited pronounced ultrastructural abnormalities including internal cavities, collapsed morphology, loss of cytoplasmic contents, incomplete exine formation characterized by missing bacula and reduced sporopollenin deposition (Fig. 2G-I). These results indicate that HDA6 is essential for proper pollen development and maturation in Arabidopsis.

### HDA6 epigenetically represses *ANAC087* to restrict *CEP1* expression during tapetal PCD

To explore the underlying mechanism by which HDA6 regulates tapetal PCD and male fertility, we performed RNA-seq analysis of developing flower buds from *axe1-4* and WT plants, and identified differentially expressed genes (DEGs) (Fig. 3A). In light of the premature tapetal PCD phenotype of *axe1-4*, we focus on upregulated DEGs associated with tapetum development and degeneration. Among the 2891 genes upregulated in *axe1-4*, *CEP1* was of particular interest since it was strongly upregulated and has a documented role in tapetal development (Fig. 3B). Previous studies showed that overexpression of *CEP1* in Arabidopsis triggers premature tapetal PCD at stage 9, closely resembling the phenotype observed in *axe1-4* (Fig. 2D) (*20*). Consistent with this possible association, qRT-PCR confirmed that *CEP1* transcript levels were significantly elevated in *axe1-4* (Fig. 3C). We therefore examined whether HDA6 directly controls *CEP1* transcription. However, dual-luciferase (dual-LUC) assays in *Nicotiana benthamiana* leaves showed that HDA6 did not directly alter *CEP1* promoter activity (Fig. 3D), and chromatin immunoprecipitation coupled with quantitative PCR (ChIP-qPCR) analysis detected no significant change in H3K9/K14Ac enrichment at the *CEP1* locus in *axe1-4* relative to WT (Fig. 3E). These findings suggest that HDA6 regulates *CEP1* expression indirectly rather than through direct transcriptional control at the *CEP1* promoter.

**Figure 3.**
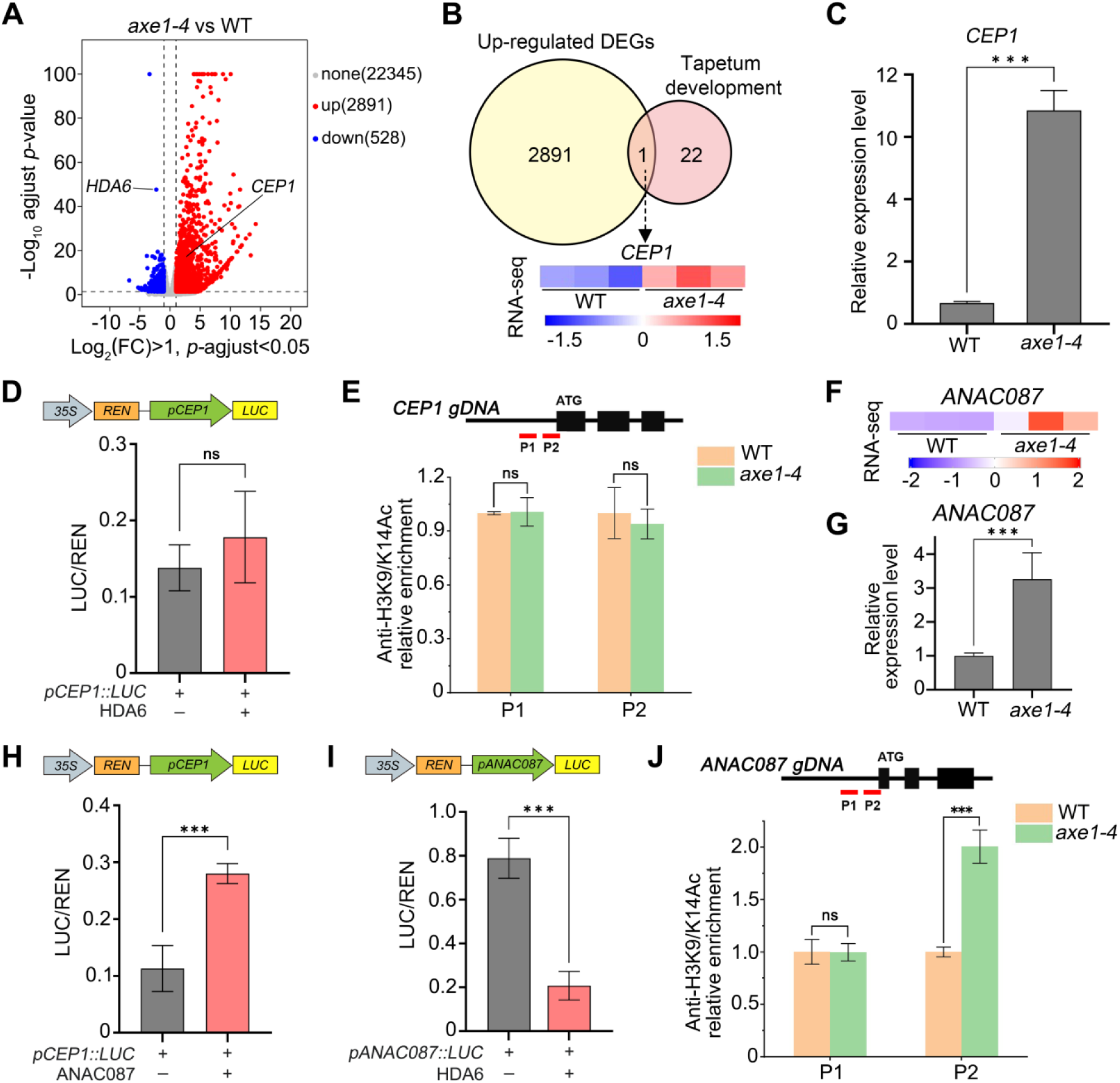
HDA6 represses *ANAC087* to indirectly restrict *CEP1* expression. (**A**) Volcano plot of differentially expressed genes (DEGs) identified by RNA-seq in early inflorescences of *axe1-4* compared to WT. Vertical dashed lines indicate log_2_ (fold change) = ±1, and the horizontal dashed line indicates *P-*adjusted = 0.05. Genes with *P-*adjusted < 0.05 and Log_2_ (fold change) > 0.5 were defined as upregulated DEGs. (**B**) Venn diagram showing the overlap between genes upregulated in *axe1-4* and genes associated with tapetum development. The lower panel shows the RNA-seq expression pattern of *CEP1* in WT and *axe1-4*. (**C**) qRT-PCR analysis of *CEP1* transcript levels in early inflorescences of WT and *axe1-4* (n = 3). (**D**) Dual-luciferase assay showing that HDA6 does not directly alter *CEP1* promoter activity in *Nicotiana benthamiana* leaves (n = 3). (**E**) ChIP-qPCR analysis of H3K9/K14 acetylation enrichment at the *CEP1* locus in early inflorescences of WT and *axe1-4* (n = 3). P1 and P2 indicate the analyzed regions. (**F**) RNA-seq expression pattern of *ANAC087* in WT and *axe1-4*. (**G**) qRT-PCR analysis of *ANAC087* transcript levels in early inflorescences of WT and *axe1-4* (n = 3). (**H**) Dual-luciferase assay showing that ANAC087 activates the *CEP1* promoter (n = 3). (**I**) Dual-luciferase assay showing repression of the *ANAC087* promoter by HDA6 (n = 3). (**J**) ChIP-qPCR analysis of H3K9/K14 acetylation enrichment at the *ANAC087* locus in early inflorescences of WT and *axe1-4* (n = 3). P1 and P2 indicate the analyzed regions. All of the samples in (**A**-**C** and **E**-**G** and **J**) were collected from early flowers before the 15^th^ flower. Values are means ±SD. ns, not significant. \**P* < 0.05, \*\**P* < 0.01, and \*\*\**P* < 0.001.

Given that HDA6 did not directly regulate *CEP1* transcription, we next asked whether it controls an upstream activator of *CEP1*. The transcription factor ANAC087 has been reported to activate *CEP1* and function in leaf senescence in Arabidopsis (*21*), while its role in tapetal PCD remains unknown. We therefore examined whether the expression level of *ANAC087* is altered in *axe1-4*. Both transcriptome profiling and qRT-PCR analysis showed that *ANAC087* expression was significantly increased in the mutant (Fig. 3F and G). To verify that ANAC087 can activate *CEP1* transcription, we performed dual-LUC assays and found that co-expression of *ANAC087* with *pCEP1::LUC* markedly enhanced LUC activity (Fig. 3H), consistent with previous findings during Arabidopsis leaf senescence (*21*). We next tested whether HDA6 regulates *ANAC087* transcription. Co-expression of *HDA6* with *pANAC087::LUC* modestly reduced promoter activity (Fig. 3I), and ChIP-qPCR analysis further showed that H3K9/K14Ac enrichment at the *ANAC087* promoter was significantly increased in *axe1-4* compared with WT (Fig. 3J). These findings demonstrate that HDA6 represses *ANAC087* transcription by reducing histone acetylation at its promoter, thereby indirectly restraining *CEP1* expression.

### SPL8 cooperates with HDA6 to repress *ANAC087* expression

Given that HDA6 is often recruited by sequence-specific transcription factors to regulate target gene expression (*28, 29*), we therefore hypothesized an upstream transcription factor might recruit HDA6 to the *ANAC087* locus during the timely onset of tapetal PCD. To identify such a factor, we performed a candidate-based screen focusing on Arabidopsis transcription factors previously implicated in tapetal PCD (*20, 25*). Luciferase complementation assays identified SPL8 as a candidate HDA6-interacting protein (Fig. 4A and B, and Fig. S6). This interaction was further validated by fluorescence resonance energy transfer (FRET) and co-immunoprecipitation (Co-IP) assays in Arabidopsis (Fig. 4C and D). Moreover, *in vitro* pull-down assays confirmed a direct interaction between HDA6 and SPL8 (Fig. 4E). Together, these results identify SPL8 as an HDA6-interacting factor that may participate in the repression of *ANAC087* transcription.

**Figure 4.**
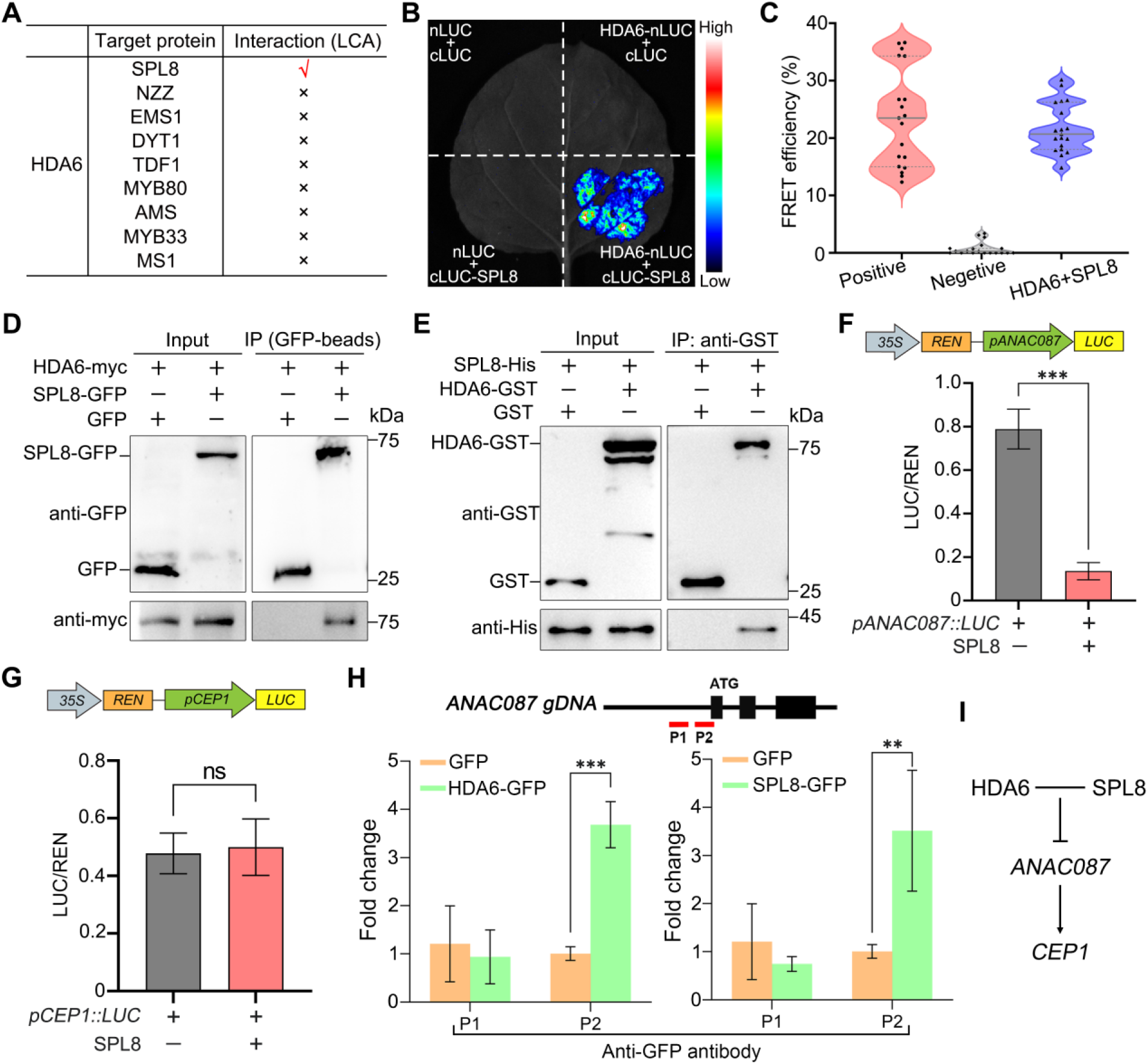
HDA6 interacts with SPL8 and cooperatively represses *ANAC087* expression. (**A**) Candidate-based LCA screen for interactions between HDA6 and transcription factors implicated in tapetal development and degeneration. SPL8 was identified as a candidate HDA6-interacting factor. (**B**) Representative images of Representative LCA images showing the interaction between HDA6 and SPL8 in *Nicotiana benthamiana* leaves (n = 4). (**C**) FRET efficiency analysis of the protein interaction between HDA6 and SPL8. CFP-YFP and free CFP plus YFP were used as positive and negative controls, respectively (n ≥ 18). (**D**) Representative results of Co-IP assay showing interaction between HDA6-myc and SPL8-GFP in transgenic Arabidopsis. (n = 4). (**E**) Representative results of *In vitro* GST pull-down assay confirming direct interaction between HDA6-GST and SPL8-His (n = 4). (**F** and **G**) Dual-luciferase assays showing that SPL8 represses the *ANAC087* promoter (**F**), while does not significantly affect the *CEP1* promoter (**G**) (n = 3). (**H**) CUT&Tag-qPCR analysis showing enrichment of HDA6-GFP and SPL8-GFP at the *ANAC087* locus (n = 3). P1 and P2 indicate the analyzed regions. All of the samples in (**H**) were collected from early flowers before the 15^th^ flower. (**I**) Brief model for cooperative repression of *ANAC087* by HDA6 and SPL8, leading to restriction of *CEP1* expression. Values are means ±SD. ns, not significant. \**P* < 0.05, \*\**P* < 0.01, and \*\*\**P* < 0.001. NZZ, nozzle; EMS1, excess microsporocyte 1; DYT1, dysfunctional tapetum 1; TDF1, defective in tapetal development and function 1; MYB80, MYB domain protein 80; AMS, aborted microspores; MYB33, MYB domain protein 33; MS1, male sterility 1.

In addition, the reported phenotype of the Arabidopsis T-DNA insertion mutant *spl8* including early floral sterility and aborted pollen closely resembles that of *axe1-4* (*25*). This phenotypic similarity suggests that SPL8 and HDA6 may act in a common pathway controlling tapetal PCD and male gametophyte development. To test whether SPL8 represses *ANAC087* transcription, we performed dual-LUC assays. Co-expression of *SPL8* with *pANAC087::LUC* significantly reduced LUC activity, whereas co-expression of *SPL8* and *pCEP1::LUC* had no significant effect (Fig. 4F and G), suggesting that SPL8 represses *ANAC087* but does not directly regulate *CEP1*. To determine whether HDA6 and SPL8 directly bind the *ANAC087* genomic region, we performed Cleavage Under Targets and Tagmentation (CUT&Tag) assays using flower buds from *pHDA6::HDA6-GFP* and *pSPL8::SPL8-GFP* transgenic lines. CUT&Tag-qPCR detected significant enrichment of both proteins at the *ANAC087* locus (Fig. 4H). These results support a model in which SPL8 cooperates with HDA6 at the *ANAC087* locus to repress *ANAC087* transcription and thereby limit *CEP1* expression (Fig. 4I). Furthermore, to assess the temporal coherence of the proposed regulatory pathway, we analyzed *HDA6*, *SPL8*, *ANAC087* and *CEP1* expression across anther and tapetal development using multiple transcriptomic datasets. *HDA6* and *SPL8* exhibited similar expression profiles, with relatively high transcript levels in early anthers (stages 4-7) and tapetum (stages 6-7), followed by a progressive decline during subsequent development. Their expression was markedly reduced during anther maturation and dehiscence, when *SPL8* transcripts became undetectable (Fig. S7A). By contrast, *ANAC087* expression was low at early stages and gradually increased toward anther maturity (Fig. S7B). *CEP1* showed a similar profile but was nearly undetectable in early anthers and tapetum. Collectively, these independent RNA-seq datasets reveal temporally coordinated expression patterns consistent with the proposed HDA6-SPL8-*ANAC087*-*CEP1* regulatory cascade (Fig. 4I and Fig. S7).

### HDA6 and SPL8 repress the *ANAC087-CEP1* pathway to safeguard early male fertility in Arabidopsis

To further assess the role of SPL8 in plant male reproductive development, we generated two independent CRISPR/Cas9 mutation alleles, *spl8-3* and *spl8-10*. Both mutants exhibited early-stage fertility defects similar to those of the *spl8* T-DNA insertion mutant and *axe1-4* (Fig. S8). In *spl8-3*, seed set gradually recovered in later flowers, resembling the temporal pattern observed in *axe1-4* (Fig. 5A and B). This similarity led us to test whether *ANAC087* and *CEP1* functionally mediate the early sterility of *axe1-4*. RNAi-mediated knockdown of either *ANAC087* or *CEP1* in the *axe1-4* background substantially rescued the early sterility phenotype and largely restored seed set in early siliques (Fig. 5A and B and Fig. S8). Consistent with these genetic results, qRT-PCR analysis showed that both *ANAC087* and *CEP1* were significantly upregulated in *spl8-3*, as in *axe1-4* (Fig. 5C).

**Figure 5.**
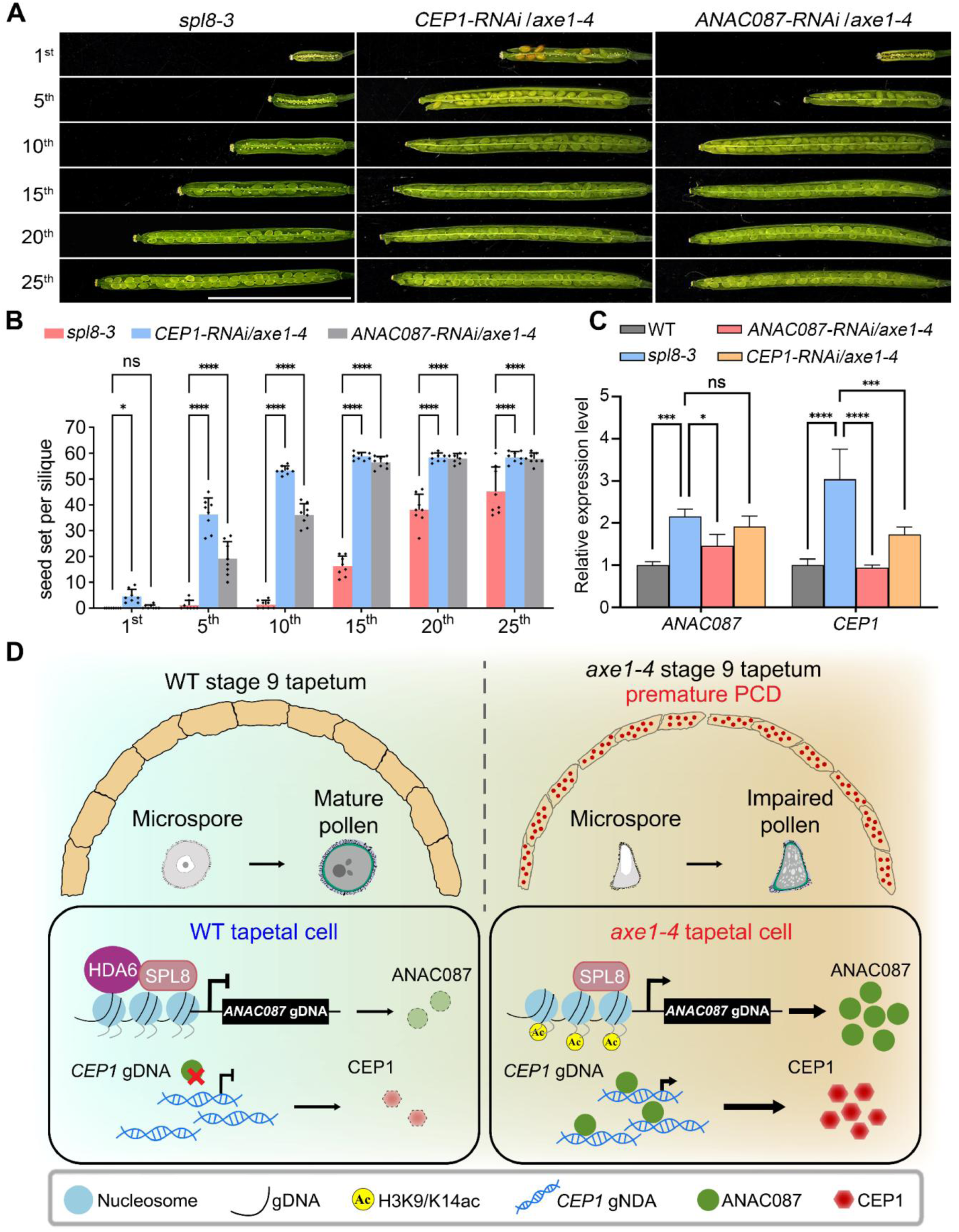
HDA6-mediated epigenetic repression of the *ANAC087*-*CEP1* pathway is required for early male fertility in Arabidopsis. (**A**) Representative siliques from the indicated flower positions in *spl8-3*, *CEP1-RNAi/axe1-4* and *ANAC087-RNAi/axe1-4.* (**B**) Quantification of seed number per silique at the indicated flower positions in *spl8-3*, *CEP1-RNAi/axe1-4* and *ANAC087-RNAi/axe1-4* plants (n = 8). (**C**) qRT-PCR analysis of *ANAC087* and *CEP1* expression level in early inflorescence of WT, *spl8-3*, *ANAC087-RNAi/axe1-4* and *CEP1-RNAi/axe1-4* (n = 3). All of the samples in (**C**) were collected from early flowers before the 15^th^ flower. (**D**) Schematic working model for the HDA6-SPL8-*ANAC087*-*CEP1* regulatory pathway controlling tapetal PCD and pollen development during early flowering. In WT anther tapetal cell at stage 9, HDA6 and SPL8 cooperatively repress *ANAC087* expression, which is associated with reduced histone acetylation at the *ANAC087* locus and restriction of downstream *CEP1* activation, thereby preventing precocious tapetal PCD and ensuring pollen maturation and fertility. In *axe1-4* mutant, loss of HDA6 results in elevated histone acetylation at the *ANAC087* locus, derepression of *ANAC087*, activation of *CEP1*, and premature tapetal PCD, ultimately leading to defective pollen development and male sterility. Scale bars = 10 μm in (**A**). Values are means ± SD. ns, not significant. \**P* < 0.05, \*\**P* < 0.01, \*\*\**P* < 0.001 and \*\*\*\**P* < 0.0001.

Thus, these data indicate that SPL8 suppresses *ANAC087*-*CEP1* pathway and further support the conclusion that derepression of this pathway underlies the early sterility phenotype of *axe1-4*. Collectively, our findings support a model in which HDA6 and SPL8 cooperatively mediate epigenetic repression of *ANAC087*, thereby restricting the *ANAC087-CEP1* pathway and ensuring proper male gametophyte development and fertility specially confined at the early flowering stage (Fig. 5D). Extending this mechanistic model beyond Arabidopsis, phylogenetic analysis showed that HDA6 orthologs are broadly conserved across various plant species but divergent from yeast ScRPD3 and mammalian HDAC1/2-type deacetylases (Fig. S9). It suggests that HDA6-mediated reproductive regulation may represent a conserved chromatin-based mechanism in plants.

## Discussion

Our study provides, to our knowledge, the first evidence that HDA6 contributes to the epigenetic regulation of pollen maturation and male fertility during the early flowering stage in Arabidopsis. Mechanistically, HDA6 acts together with SPL8 to repress *ANAC087* through histone deacetylation, thereby restricting activation of the ANAC087-CEP1 cell death pathway and preventing premature tapetal PCD (Fig. 5). This HDA6-dependent chromatin regulatory machinery is therefore essential for maintaining the proper transcriptional state required for timely tapetal degeneration and pollen maturation in early flowers. A notable feature of the *hda6* mutant is their dynamic and stage-dependent fertility phenotype (Fig. 1). Seed number per silique is severely reduced during early reproductive development but gradually recovers to near-normal levels at middle to late flowering stages (Fig. 1). This early sensitivity followed by late recovery phenotypic pattern distinguishes *hda6* from completely sterile mutants and indicates that HDA6-SPL8-mediated epigenetic regulation is particularly critical within a defined developmental window (*20, 30*). Our findings suggest that male reproductive development is not controlled by a single uniform regulatory program throughout the flowering period. Instead, pollen maturation and fertility may depend on at least two temporally distinct regulatory systems, with an HDA6-dependent epigenetic pathway functioning during early flowering and a separate or compensatory regulatory machinery supporting male reproductive development at later stages.

Nevertheless, this temporal recovery also raises important questions regarding developmental compensation and reproductive plasticity. In natural environments, plants must balance reproductive output with resource availability, and allocation trade-offs between early and late reproductive structures are central to plant life-history strategies (*31, 32*). In this context, the HDA6-SPL8 module may function as a developmental checkpoint that safeguards the precision of chromatin-mediated transcriptional regulation during the vulnerable transition from vegetative to reproductive growth. When this checkpoint is compromised, early flowers are highly susceptible to premature tapetal degeneration and defective pollen maturation. However, the later restoration of fertility implies that additional regulatory pathways, developmental cues or resource-dependent compensatory mechanisms can partially bypass or buffer the loss of HDA6-SPL8 function. Thus, our findings define the HDA6-SPL8-mediated repression of the *ANAC087-CEP1* pathway as an essential epigenetic checkpoint for early anther development (Fig. 5). At the same time, identifying the late-acting mechanisms that restore pollen maturation and male fertility will be crucial for understanding how plants coordinate reproductive robustness, developmental plasticity and fertility across the flowering period.

Considered in light of previous studies, our findings introduce an epigenetic dimension to the current framework of tapetum regulation which has been shaped largely by transcription factor networks that control anther development such as the DYT1-TDF1-AMS-MS188-MS1 cascade (*30, 33–36*). Although these pathways are central to tapetal development, degeneration and pollen maturation, the contribution of chromatin-based regulation to tapetal PCD remains less clearly defined. Existing evidence for epigenetic regulation in male reproductive development has been derived primarily from studies of siRNA-mediated pathways, whereas the possible involvement of histone modification in tapetal degeneration has received comparatively less attention (*37*). Our findings suggest that histone deacetylation constitutes an additional regulatory layer contributing to the proper progression of tapetal PCD and pollen development. In addition, we also find that HDA6 affects ovule numbers without measurably impairing female gametophytic fertility (Fig. 1 and Fig. S3), suggesting a broader role for HDA6 in the regulation of plant sexual reproduction, although the underlying mechanism remains to be elucidated. This view is consistent with the well-established functions of HDA6 as a pleiotropic epigenetic regulator in RNA-directed DNA methylation, transposon silencing, flowering-time regulation and various stress responses (*38–40*). The restriction of the fertility defect largely to early flowering further suggests that HDA6 activity may be particularly important during a narrow developmental window, possibly through interaction with stage- and tissue-specific transcription factors such as SPL8 (Fig. 4). In addition, the confinement of the phenotype to early flowering raises the possibility that male reproductive development is coordinated by distinct regulatory machineries operating at different stages of the flowering period.

In fact, the functions of HDA6 in plant development and stress responses have been extensively characterized, yet its contribution to male fertility has remained largely unrecognized (*41*). One possible explanation for this gap is the predominant use of the *axe1-5* allele in previous HDA6 studies, whereas the *axe1-4* allele has been examined far less frequently. EMS-induced mutagenesis of *HDA6* generated a series of independent alleles, including *axe1-1* to *axe1-5*, *sil1*, and *shi5*, each carrying distinct mutations and exhibiting different degrees of phenotypic severity (*42*). This allelic diversity suggests that the developmental consequences of HDA6 dysfunction are strongly influenced by the position and nature of the mutation. Indeed, *axe1-5*, the most widely used allele, has been associated with altered jasmonate responsiveness, delayed leaf senescence, changes in flowering time and relatively mild leaf morphological defects, but it does not display obvious fertility defects during the early reproductive stage (*14, 43*). By contrast, *axe1-4* exhibits more pronounced developmental abnormalities including severe defects in leaf morphogenesis, and as shown in this study, a marked reduction in male fertility during early reproductive development (Fig. 1 and 2) (*16*). The phenotypic divergence between these alleles is likely related to their distinct mutational sites within *HDA6*. Previous sequencing analyses showed that the *axe1-4* mutation is located within the histone deacetylase catalytic domain, whereas the *axe1-5* mutation lies near the C terminus and outside this domain (*42, 44*). Consistently, CRISPR/Cas9-generated *hda6* mutant lines phenocopy the severe early male fertility defects observed in *axe1-4*, supporting the view that disruption of the deacetylase domain causes a stronger impairment of HDA6 function (Fig. 1). Therefore, the male fertility phenotype observed in *axe1-4*, but not in *axe1-5*, is likely attributable to allele-specific differences in the extent of HDA6 functional disruption. These findings emphasize the importance of allelic context when interpreting HDA6 function and suggest that the use of the *axe1-4* allele uncovered a previously overlooked epigenetic role of HDA6 in regulating male reproductive development.

In addition, phylogenetic analysis showed that *HDA6* orthologs are widely distributed across angiosperms, including both dicots and monocots, and form a plant HDA6 lineage within the conserved RPD3/HDA1-type class I HDAC family, distinct from yeast ScRPD3 and mammalian HDAC1/2 (Fig. S9). This phylogenetic separation suggests lineage-specific diversification of class I HDACs in plants, consistent with the involvement of Arabidopsis HDA6 in RdDM-associated heterochromatin silencing, transposable element silencing and developmental phase transition (*36*). By contrast, mammalian HDAC1/2 commonly function as catalytic components of large chromatin-regulatory complexes such as switch-independent 3 (Sin3), nucleosome remodeling and deacetylase (NuRD) and RE1-silencing transcription factor corepressor (CoREST), to regulate cell-cycle progression, differentiation, DNA repair and genome stability (*38, 45*).

Despite these differences, our findings suggest a possible conserved biological principle in reproductive development. We show that HDA6 contributes to male fertility in Arabidopsis by regulating the timing of tapetal PCD, thereby supporting pollen maturation (Fig. 2). In mammals, HDAC1 has also been implicated in male reproduction. It interacts with bromodomain testis-specific protein (BRDT) which is a member of the bromodomain and extra-terminal domain (BET) family to repress *H1t* transcription during spermatogenesis, thereby contributing to the control of male fertility (*46*). Although plant and animal male reproductive systems differ substantially in cellular organization, developmental context and reproductive structures, it suggests that RPD3/HDA1-type histone deacetylases may share a broadly conserved role in coordinating transcriptional timing during male reproductive development. In this context, HDA6-mediated repression in the tapetum and HDAC1-mediated regulation in spermatogenic cells may represent divergent implementations of a common regulatory strategy, in which chromatin-modifying enzymes cooperate with partner proteins to fine-tune gene expression that required for gametophyte or germ-cell maturation.

## Materials and Methods

### Plant materials and growth conditions

*Arabidopsis thaliana* plants were grown at 22°C under long-day (LD) conditions with 16 h light/8 h dark cycle photoperiod. Seeds were stratified at 4°C for 2 days and then placed on 1/2 Murashige and Skoog (MS) medium supplemented with 1% sucrose and 1% agar (pH 5.7). All plants used in this study were in the Columbia (Col-0) background. Transgenic lines were generated by the floral dip method (*47*). The *axe1-4* mutant, *pHDA6::HDA6-GFP/axe1-4* and *pHDA6::HDA6-myc/axe1-4* lines were used as previously described (*16, 44*). Transgenic Arabidopsis lines expressing *pUBQ::GFP* and *pSPL8::SPL8-GFP* were generated by transforming Col-0 plants with the corresponding constructs. *ANAC087*-*RNAi*/*axe1-4* and *CEP1*-*RNAi*/*axe1-4* transgenic Arabidopsis were generated by introducing *ANAC087*-RNAi and *CEP1*-RNAi constructs into the *axe1-4* background respectively. All *hda6* and *spl8* mutant alleles were generated using the previously reported CRISPR/Cas9 system (*48*). Gene-specific guide RNAs targeting *HDA6* and *SPL8* genes were designed using CRISPR-GE (*49*). Fragment containing two sgRNAs for *HDA6* and *SPL8* were amplified from *pCBC-DT1T2* and cloned into *pHEE401E* by Golden Gate cloning to generate the constructs *pHEE401E-HDA6* and *pHEE401E-SPL8*. These constructs were transformed into the Col-0 plants, and transgenic lines were subsequently screened for targeted mutations. The full list of PCR primers is provided in Dataset S3.

### Pollen wax extraction and GC-MS analysis

Freshly opened flowers were harvested, immediately frozen in liquid nitrogen, and stored at -80°C until analysis. Mature pollen grains were released in ddH_2_O by vortexing, and floral petals and anthers were subsequently removed with forceps. Arabidopsis pollen was pelleted by centrifugation at 800 × g for 1 min. Each biological replicate contained 100 mg of pollen, and three independent biological replicates were analyzed. After removal of the supernatant, wax extraction was carried out as previously described (*50*). Each sample was transferred to a Teflon-lined screw-cap glass tube, and 3 mL of 2.5% (v/v) sulfuric acid in methanol containing 0.2 mg/mL methyl heptadecanoate as an internal standard was added. Samples were incubated at 80°C water bath for 1 h, cooled to room temperature (RT) and then mixed with 1.5 mL pentane and 4.5 mL 0.9% (w/v) NaCl. After vigorous vortexing for 1 min, samples were centrifuged at 1200 × g for 2 min. The supernatant was transferred to a new brown glass tube and evaporated to dryness in a fume hood. The organic phase was transferred to a fresh amber glass tube and evaporated to dryness in a fume hood. The residue was resuspended in 300 μL pentane, and 150 μL was transferred to an amber glass vial fitted with an insert for analysis. Wax components were identified by their relative retention times and characteristic mass spectra (TRACE 1300; Thermo Fischer Scientific).

### mRNA-seq

Total RNA was isolated using TRIzol (15596026CN; Invitrogen), and RNA concentration, purity, and integrity were evaluated using a NanoDrop spectrophotometer (NDULTRAGL; Thermo Fischer Scientific). For each sample, 3 μg of total RNA was used for library construction. mRNA was purified using poly(T) oligo-attached magnetic beads and fragmented in Illumina proprietary fragmentation buffer under elevated temperature. First-strand cDNA was synthesized using random oligonucleotides and SuperScript II, followed by second-strand synthesis using DNA Polymerase I and RNase H. After end repair and 3’ adenylation, Illumina paired-end adapters were ligated to the cDNA fragments. Sequencing libraries were generated using NEB Next UltraTM RNA Library Prep Kit for Illumina (E7770; NEB) following manufacturer’s recommendations, and index codes were added to attribute sequences to each sample. The library preparations were sequenced on an Illumina Hiseq X Ten platform and paired-end reads were generated. The RNA-seq clean reads mapped to the Arabidopsis genome (TAIR10) using TopHat (v2.1.1). DEseq (v1.10.1) was applied for differential gene expression analysis. Genes with q ≤ 0.05 and |log2_ratio| ≥ 1.5 were identified as DEGs.

### *In vitro* and *in vivo* Arabidopsis pollen germination

Both *in vitro* and *in vivo* Arabidopsis pollen germination are performed as previously described (*51–53*). Briefly, for *in vitro* pollen germination, pollen grains from newly opened flowers were spread onto solid pollen germination medium composed of 18% sucrose, 0.01% boric acid, 5 mM CaCl_2_, 5 mM KCl, 1 mM MgSO_4_, and 0.5% agarose (pH 7.5). The plates were placed upside down in a humid chamber and incubated at 28°C for 3 h. Germinated pollens were subsequently examined and imaged using a stereomicroscope (M205 FA; Leica). For *in vivo* pollen germination, flower buds with visible white petal tips were selected and emasculated. Freshly opened flowers were then used as pollen donors, and pollen grains were gently transferred from the stamens onto recipient pistils. At 10 h after pollination, pistils were excised and fixed in freshly prepared Carnoy’s fixative (Ethanol : Acetate = 3:1, v/v) for 2 h at RT. After fixation, pistils were rehydrated through a graded ethanol series of 70%, 50%, and 30% (v/v) followed by ddH_2_O, with each step performed for 5 min. Samples were then cleared in 6 M NaOH at RT for 12 h. After clearing, the pistils were washed five to six times with ddH_2_O and stained with freshly prepared 0.2% aniline blue in the dark for 2 h. Pollen germination and pollen tube growth were subsequently observed and imaged under a fluorescence microscope (DM6 B; Lecia) using UV excitation.

### Sample treatment and scanning electron microscopy

For pollen exine staining, freshly collected pollen grains were incubated with 30 μL of 0.1% (w: v) Auramine O solution. After gentle resuspension, samples were incubated in the dark for 3 min and subsequently observed and imaged using a confocal laser scanning microscope (TCS SP8; Lecia). For DAPI staining, mature Arabidopsis pollen grains were collected and concentrated by centrifugation at 800 × g for 1 min. After removal of the supernatant, the pollen pellet was gently resuspended in 20 μL of DAPI staining solution containing 0.25 M sodium phosphate (pH 7.0), 0.25 mM EDTA, 0.025% Triton X-100 (v/v) and 0.1 µg/ml DAPI. Samples were incubated in the dark for 30 to 45 min and immediately imaged using a confocal laser scanning microscope (TCS SP8; Leica). For scanning electron microscopy, mature pollen grains were collected from freshly dehisced anthers and mounted on scanning electron microscopy stubs. The samples were sputter-coated with palladium-gold using a sputter coater (EM-ACE600; Leica) and observed by scanning electron microscopy (EVO MA15, Zeiss) at an acceleration voltage of 10 kV.

### RNA *in situ* hybridization

mRNA *in situ* hybridization was performed to examine the spatial expression patterns of genes of interest, following a standard protocol (*54*). To generate probes, gene-specific DNA fragments were amplified from cDNA using primers containing T3 or T7 promoter sequences. Fragments of ∼500 bp, corresponding to variable coding regions and untranslated regions (UTRs), were used as templates for *in vitro* transcription. Digoxigenin-labeled sense and antisense RNA probes were synthesized using T3 RNA polymerase (11031163001; Roche), T7 RNA polymerase (10881775001; Roche) and digoxigenin-11-UTP (11209256910; Roche). Color development was performed until optimal signal intensity was obtained, typically for approximately 3 d.

### Semithin sections and transmission electron microscopy

Semithin sectioning was performed essentially as described in previous studies (*55*). Intact Arabidopsis inflorescences were collected and open flowers were removed before fixation. Samples were fixed in 4% glutaraldehyde under vacuum for 30 min and then incubated overnight at 4°C. Following primary fixation, samples were washed six times with phosphate-buffered saline (PBS) with each wash lasting 15 min. Samples were then post-fixed in 2% osmium tetroxide for 2 to 3 h at RT in the dark. After postfixation, samples were rinsed six times with distilled water, 15 min per wash, and dehydrated through a graded acetone series of 30%, 40%, 50%, 60%, 75%, 85%, and 95% (v/v) followed by three changes of 100% acetone for 10 min each. Dehydrated samples were infiltrated sequentially with acetone/resin mixtures at ratios of 5:1, 3:1, 1:1, 1:3, and 1:5 (v/v) followed by pure resin, with each step carried out for 12 h. The resin was replaced at least twice before embedding. Samples were then transferred to embedding molds containing fresh resin and properly oriented prior to polymerization at 65°C for 2 to 3 d. Semithin sections of 1 μm thickness were prepared using a microtome (RM2155, Leica) equipped with a glass knife (7890-04; Leica). Sections were stained with 0.1% (w/v) toluidine blue and examined under a light microscope. For ultrathin sections, ultrathin sections with a thickness of 60 to 70 nm were prepared using an ultramicrotome (ARTOS 3D; Leica) equipped with a diamond knife and subsequently imaged using a transmission electron microscope (Talos L120C; FEI).

### TUNEL assay

TUNEL assays was conducted as previously described (*20*). In general, intact Arabidopsis inflorescences were collected and open flowers were removed before fixation. Samples were fixed in PBS buffer containing 4% (w/v) paraformaldehyde under vacuum infiltration for 30 min and then maintained in the same fixation buffer overnight at 4°C. Fixed flower buds were embedded and 8-μm sections were prepared for TUNEL analysis. DNA fragmentation was detected using the DeadEnd Fluorometric TUNEL System (G3250; Promega) according to the manufacturer’s protocol. Samples were examined and imaged using a fluorescence microscope (DM6 B; Leica).

### Luciferase complementation assay

Luciferase complementation assay was performed as previously described (*55, 56*). For luciferase complementation assays, coding sequence of target genes were cloned into the *pCAMBIA-35S-nLUC* and *pCAMBIA-35S-cLUC* vectors respectively. The resulting plasmids were transformed into *Agrobacterium tumefaciens* GV3101 and infiltrated into *Nicotiana benthamiana* leaves using a syringe. The tobacco plants were placed in darkness for 24 h and then incubated in a 16-h light /8-h dark photoperiod for an additional 24 h. Thereafter, the leaves were sprayed with a 1 mM luciferin solution and then kept in darkness for 5 min to quench the fluorescence. A deep cooling charge-coupled device imaging apparatus (5200 Multi; Tanon) was used to capture fluorescence images.

### FRET assays

FRET assays was performed as previously described (*55, 56*). For the FRET assay, the constructs including *pUBQ10::HDA6-CFP, pUBQ10::SPL8-YFP, pUBQ10::CFP, pUBQ10::YFP and pUBQ10::CFP-YFP* were transiently coexpressed in tobacco BY-2 protoplasts. After incubation for 10-12 h at 28°C in darkness, FRET analysis was performed using a Leica TCS SP8 confocal microscopy system according to the manufacturer’s instructions. CFP and YFP were excited at 405 and 514 nm, respectively. The CFP-YFP fusion construct which contains a short linker between CFP and YFP, served as the positive control, whereas the coexpression of unfused CFP and YFP proteins was used as the negative control. Protoplasts expressing the indicated CFP- and YFP-tagged proteins were subjected to acceptor photobleaching with a 514-nm laser at full power. FRET efficiency was calculated using the formula FRET_eff_ = (*D*_post_ − *D*_pre_)/D_post_, where *D*_post_ and *D*_pre_ represent the CFP fluorescence intensity before and after acceptor photobleaching respectively.

### Co-Immunoprecipitation

Co-immunoprecipitation assays were performed as described previously (*55, 57*). Briefly, 1 g of 7-d-old transgenic Arabidopsis flower buds expressing *pUBQ::GFP* and *pHDA6::HDA6-myc*, or *pSPL8::SPL8-GFP* and *pHDA6::HDA6-myc* was harvested and homogenized in liquid nitrogen. Total proteins were extracted in ice-cold IP buffer containing 50 mM of Tris-HCl (pH 7.4), 150 mM of NaCl, 1 mM of EDTA, 0.5% NP-40 (v/v), 10% glycerol (v/v) and 1x protease inhibitor cocktail (5892970001; Roche). After clarification by centrifugation at 16,000 × g for 15 min at 4°C, the extracts were filtered through a 0.2-μm membrane and incubated with GFP-Trap magarose beads (SM038001; Smart-Lifesciences) at 4 °C for 4 h with gentle end-to-end rotation. After five washes with ice-cold wash buffer containing 50 mM Tris-HCl (pH 7.4), 150 mM NaCl, 1 mM EDTA, 10% glycerol (v/v), bound proteins were eluted in SDS sample buffer, resolved by SDS-PAGE and subjected to immunoblot analysis with anti-GFP (A11122; Invitrogen) and anti-MYC (D191042; Sangon Biotech) antibodies. The chemiluminescence was imaged using an image analyzer (5200 Multi; Tanon).

### GST pull-down assays

GST pull-down assays were performed as previously described (*58*) with minor modifications. Generally, bait proteins were incubated with Glutathione magarose beads (SM002001; Smart-Lifesciences), and the beads were then blocked overnight in 5% BSA. After washing, bead-bound bait proteins were incubated with prey proteins for 2 h at 4°C. The beads were subsequently washed five times with buffer containing 15 mM NaCl, 20 mM Tris-HCl (pH 8.0), 0.5 mM EDTA, 0.5% NP-40 and protease inhibitor cocktail (5892970001; Roche), and the protein-bounded beads were analyzed by pull-down assay.

### Dual-luciferase reporter assay

Dual-luciferase reporter assays were performed as previously described (*21, 59*). In brief, approximately 2-kb DNA promoter sequences were cloned into the reporter vector *pGreenII::0800-LUC*, whereas the full-length coding sequences of the indicated genes were inserted into the vector *pGreenII::62-SK* as the effector constructs. The empty *pGreenII::62-SK* vector was used as a negative control. Reporter and effector plasmids were introduced into Agrobacterium strain GV3101 and coin filtrated into *Nicotiana benthamiana* leaves in the appropriate combinations. After 2 d, absolute firefly luciferase (LUC) and Renilla luciferase (REN) activities were measured using a Dual Luciferase Reporter Gene Assay Kit (11402ES60; Yeasen) on a multimode microplate reader (Spark; Tecan) according to the manufacturer’s instructions. Promoter activity was calculated as the ratio of LUC to REN.

### ChIP assay and qPCR

ChIP assays and qRT-PCR analysis were performed as previously described (*59, 60*). Developing Arabidopsis flower buds were collected, immediately frozen in liquid nitrogen, and stored at -80°C. Samples were ground into a fine powder in liquid nitrogen and chromatin was cross-linked in 1% formaldehyde. The cross-linked chromatin was isolated and sonicated to generate DNA fragments with an average size of 200 to 500 bp. Immunoprecipitation was carried out using an anti-acetyl-histone H3K9K14 antibody (06-599; Millipore) (*17*). Following recovery of the precipitated DNA, enrichment of the indicated *ANAC087* and *CEP1* regions was determined by qPCR with gene-specific primers. ChIP enrichment in *axe1-4* compared to WT was normalized to *ACTIN2* and expressed as percentage of input.

### CUT&Tag assay

CUT&Tag assays were performed as previously described with a commercial kit (TD904-01; Vazyme) (*59, 61*). Fresh young flower buds were harvested, frozen in liquid nitrogen, and ground to fine powder. Nuclei were isolated with ice lysis buffer provided with the kit, passed through a 40 µm strainer, collected by low-speed centrifugation, and further purified by density-gradient centrifugation. Purified intact nuclei were immobilized on activated ConA coated magnetic beads and permeabilized with digitonin. The bead nuclei complexes were incubated with GFP antibody (A11122; Invitrogen) overnight at 4 °C, followed by incubation with secondary antibody for 1-2 h at RT. After washing, pA/G-Tn5 transposome was added and incubated for 1 h at RT, and tagmentation was carried out in Mg^2+^-containing ChiTag buffer at 37 °C for 1 h. DNA fragments released from the tagmented chromatin were purified and amplified by PCR with indexed primers for 12 to 18 cycles. For targeted validation, enriched DNA fragments were quantified by qPCR using gene-specific primers. Each assay was performed with three biological replicates.

### Statistical analysis

Statistical analyses were performed using GraphPad Prism software (version 9.5). Differences between two groups were evaluated using Independent Student’s *t* tests. Comparisons among three or more groups were performed using one-way analysis of variance (ANOVA). Differences were considered statistically significant at *P* < 0.05.

## Accession numbers

The locus identifiers for the genes mentioned in this article are *HDA6* (AT5G63110), *CEP1* (AT5G50260), *ANAC087* (AT5G18270), *SPL8* (AT1G02065), *NZZ* (AT4G27330), *EMS1* (AT5G07280), *DYT1* (AT4G21330), *TDF1* (AT3G28470), *MYB80* (AT5G56110), *AMS* (AT2G16910), *MYB33* (AT5G06100) and *MS1* (AT5G22260).

## Acknowledgments

We apologize to those whose work could not be cited because of space restrictions. We would like to thank Prof. Shunong Bai (Peking University) and Prof. Ming Luo (South China Botanical Garden, Chinese Academy of Sciences) for providing *axe1-4* mutant, *pHDA6::HDA6-GFP/axe1-4* and *pHDA6::HDA6-myc/axe1-4* lines. We also thank members of Wang laboratory for the insightful discussion of the manuscript.

## Funding

This work is was supported by grants from the National Natural Science Foundation of China (32270358, 92354302, 91954110 and 31770196), the Natural Science Foundation of Guangdong Province (2021A1515012066) and Guangdong Provincial Specific University Discipline Construction Project (2023B10564004) to H.W.

## Author contributions

Conceptualization: K.H. and H.W. Investigation: K.H., X.W., L.Z., and J.C. Technical support: J.C. and L.Z. Analysis: K.H., C.D., H.Y. and H.W. Software: K.H. and H.W. Project administration: H.W. Funding acquisition: H.W. Supervision: H.W. Writing-original draft: K.H. Writing-review and editing: K.H. and H.W. All authors approved the final version of the manuscript for submission.

## Competing interests

The authors declare that they have no competing interests.

## Data and materials availability

All data needed to evaluate the conclusions in the paper are present in the paper and/or the Supplementary Materials. All high-throughput sequencing data of RNA-seq in this study have been deposited in NCBI BioProject database (sequence read archive database ID: PRJNA1423421).

## Notes

### Competing Interest Statement

The authors have declared no competing interest.

