## Supplemental Materials for "HDA6-mediated histone deacetylation restricts premature tapetal PCD to maintain male fertility during early flowering in Arabidopsis"

Kangwei Hu *et al.*

**This PDF file includes:**

Figs. S1 to S9

Supplementary Figures
Fig. S1

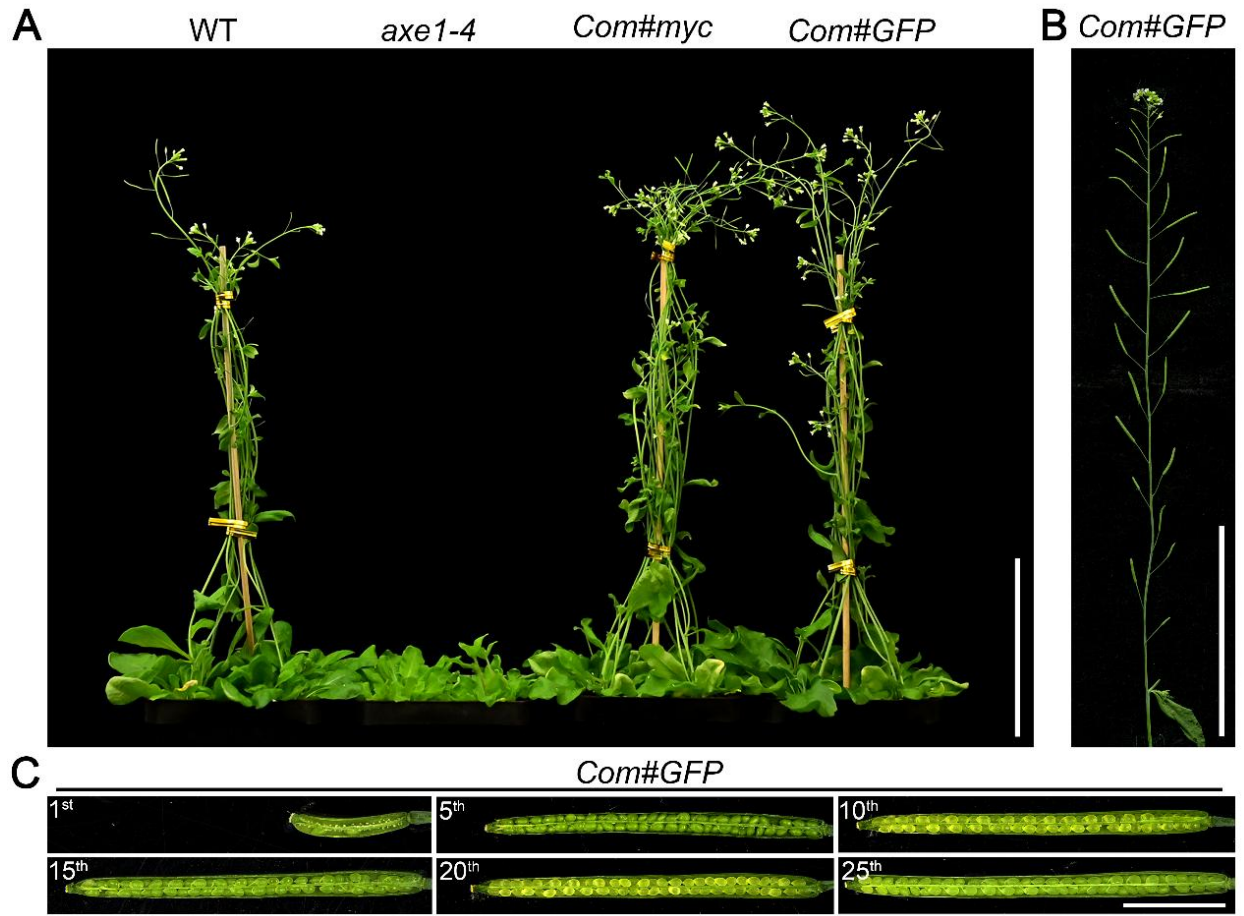

**Fig. S1. HDA6 complementation restores flowering progression and seed setting in *axe1-4*.** (A) Representative whole-plant images of WT, *axe1-4* and two independent HDA6 complementation lines at 40 d after germination. Expression of *pHDA6::HDA6-myc* or *pHDA6::HDA6-GFP* in the *axe1-4* background restored flowering progression, indicating that the delayed-flowering phenotype of *axe1-4* results from impaired HDA6 function. (B) Representative image of the primary inflorescence of the *pHDA6::HDA6-GFP* complementation line, excised below the last cauline leaf. (C) Representative images of siliques derived from the 1<sup>st</sup>, 5<sup>th</sup>, 10<sup>th</sup>, 15<sup>th</sup>, 20<sup>th</sup> and 25<sup>th</sup> flowers on the primary inflorescences of the *pHDA6::HDA6-GFP* complementation line. Scale bars = 8 cm in (A and B) and 5 mm in (C).

**Fig. S2**

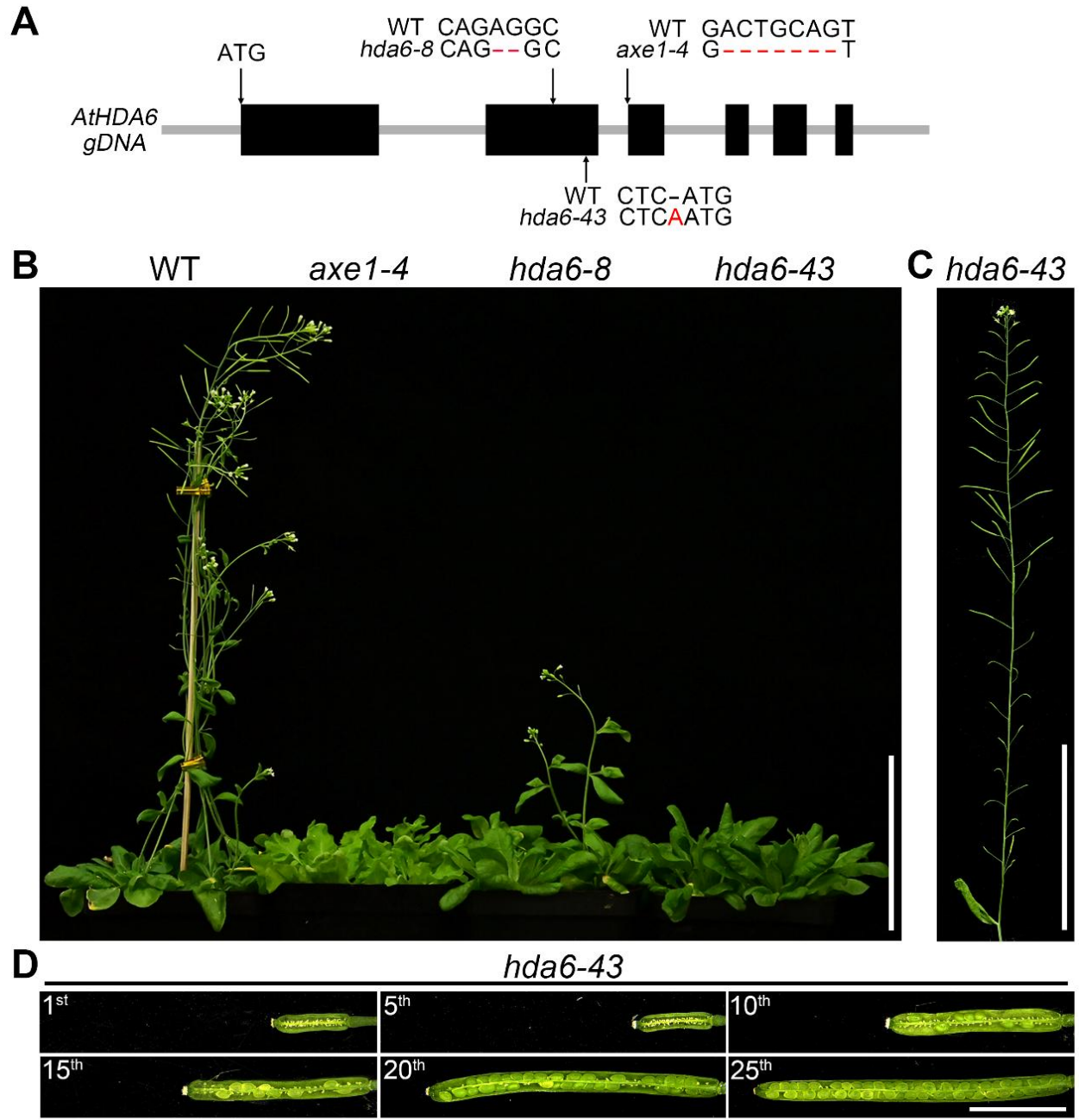

**Fig. S2. CRISPR/Cas9-generated *hda6* independent alleles recapitulate the flowering and seed setting phenotypes of *axe1-4*.**

(A) Schematic representation of the *AtHDA6* genomic locus showing the mutation sites in *axe1-4* and the CRISPR/Cas9-induced mutations in the independent alleles *hda6-8* and *hda6-43*. (B) Representative whole-plant images of WT, *axe1-4*, *hda6-8* and *hda6-43* at 40 d after germination. The delayed-flowering phenotype observed in *axe1-4* was similarly reproduced in the independent *hda6* mutant alleles. (C) Representative images of primary inflorescence of *hda6-43*, excised below the last cauline leaf. (D) Representative images of siliques derived from the 1<sup>st</sup>, 5<sup>th</sup>, 10<sup>th</sup>, 15<sup>th</sup>, 20<sup>th</sup> and 25<sup>th</sup> flowers on the primary inflorescences of *hda6-43*. Scale bars = 8 cm in (B and C) and 5 mm in (D).

Fig. S3

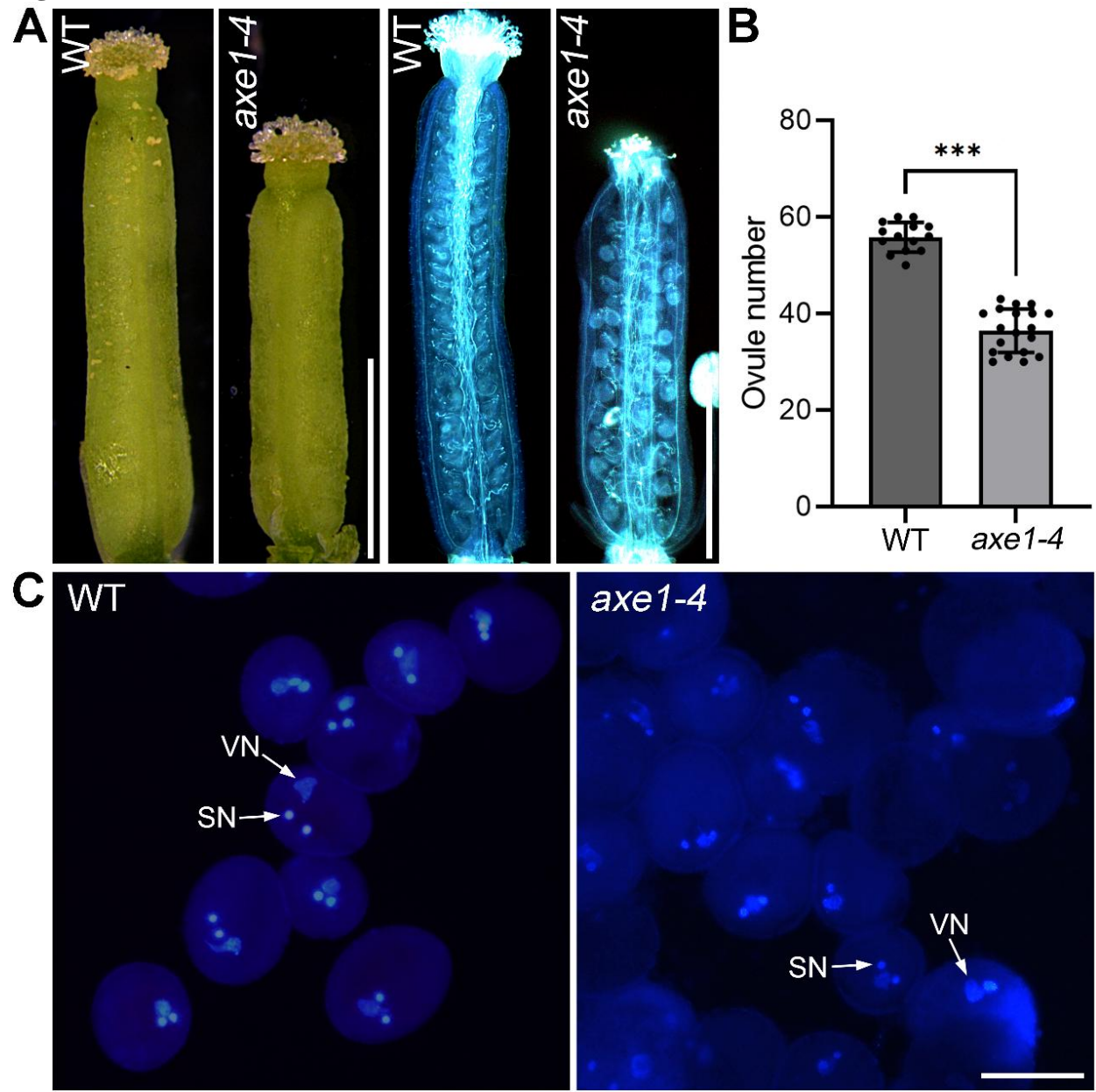

**Fig. S3. Female gametophytic function and pollen nuclear division are largely maintained in *axe1-4* despite reduced ovule number.**

(A) Representative WT and *axe1-4* pistils and aniline blue staining of *in vivo* pollen tube growth and fertilization. (B) Statistical calculation of ovule number in WT and *axe1-4* ($n \geq 14$). (C) DAPI staining of mature pollen grains of WT and *axe1-4*. All of the samples in (A-C) were collected from early flowers before the 15<sup>th</sup> flower. VN: vegetative nucleus; SN: sperm cell nucleus. Scale bars = 1 mm in (A) and 20 μm in (C). Error bars represent ± SD. \*\*\*$P < 0.001$.

Fig. S4

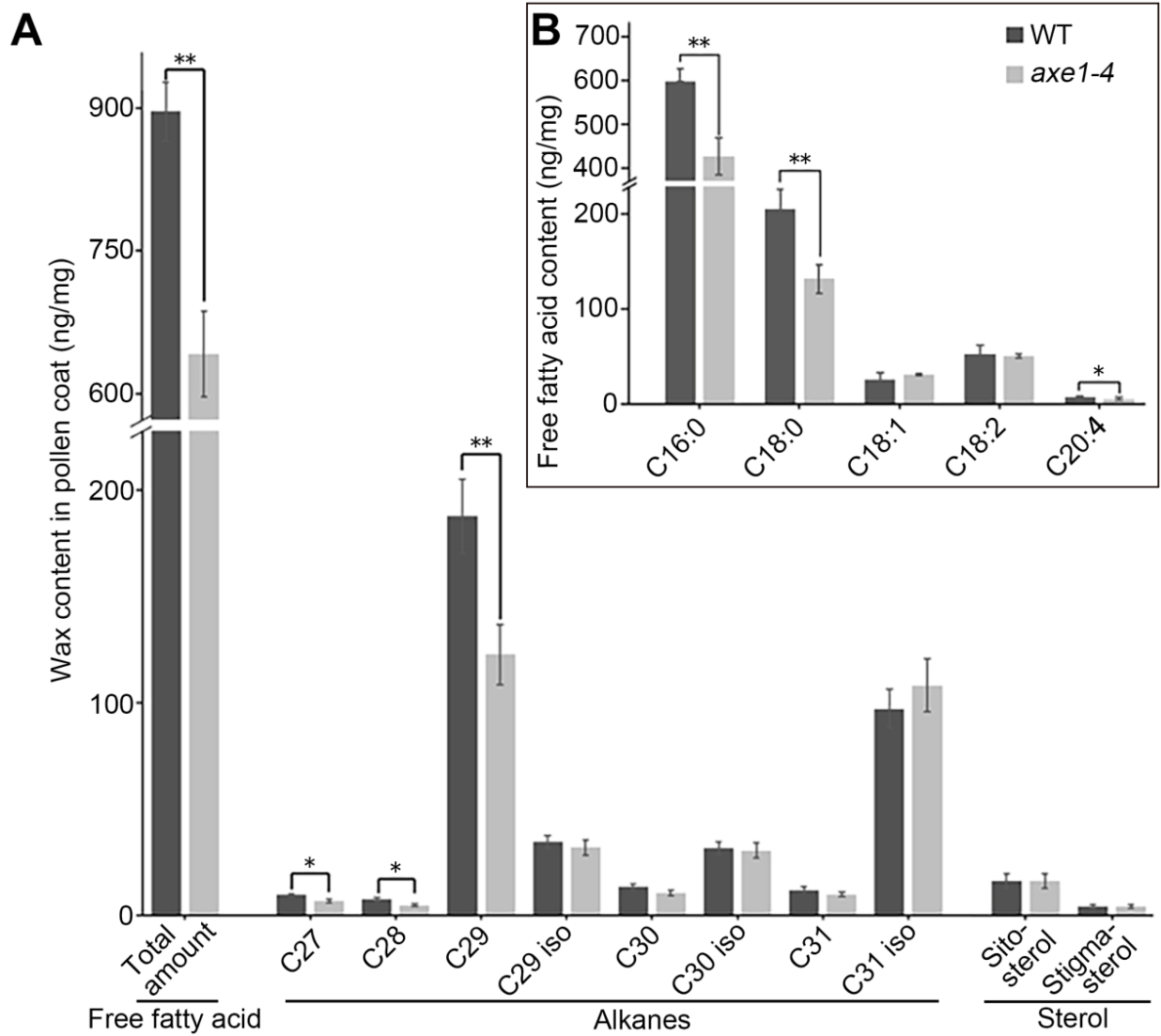

**Fig. S4. Loss of HDA6 alters pollen coat lipid accumulation in *axe1-4*.**

GC-MS-based quantification of pollen coat lipid constituents in mature pollen from WT and *axe1-4*. **(A)** Total pollen coat wax content and major long-chain lipid species including alkanes and sterols ($n = 3$). **(B)** Free fatty acid composition of mature pollens of WT and *axe1-4* ($n = 3$). Error bars represent ± SD. \*$P < 0.05$, \*\*$P < 0.01$.

**Fig. S5**

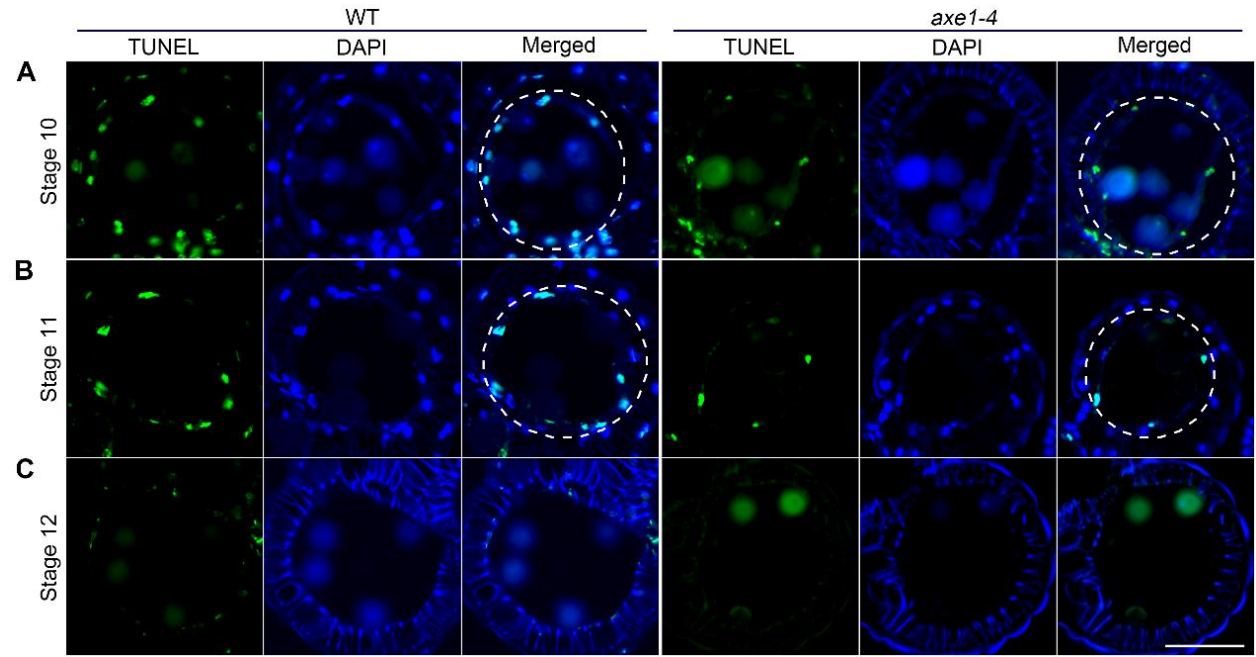

**Fig. S5. TUNEL analysis of tapetal PCD in WT and *axe1-4* mutant anthers.**

Representative transverse anther sections from WT and *axe1-4* at stages 10 to 12 were subjected to TUNEL staining to detect DNA fragmentation associated with PCD. DAPI staining marks nuclei, and merged images show the spatial relationship between TUNEL signals and anther tissues ($n \geq 3$). (A) Stage 10 anthers. (B) Stage 11 anthers. (C) Stage 12 anthers. White dashed lines indicate the anther tapetal cell layer. All of the samples in (A-C) were collected from early flowers before the 15<sup>th</sup> flower. Scale bar = 50 $\mu\text{m}$.

**Fig. S6**

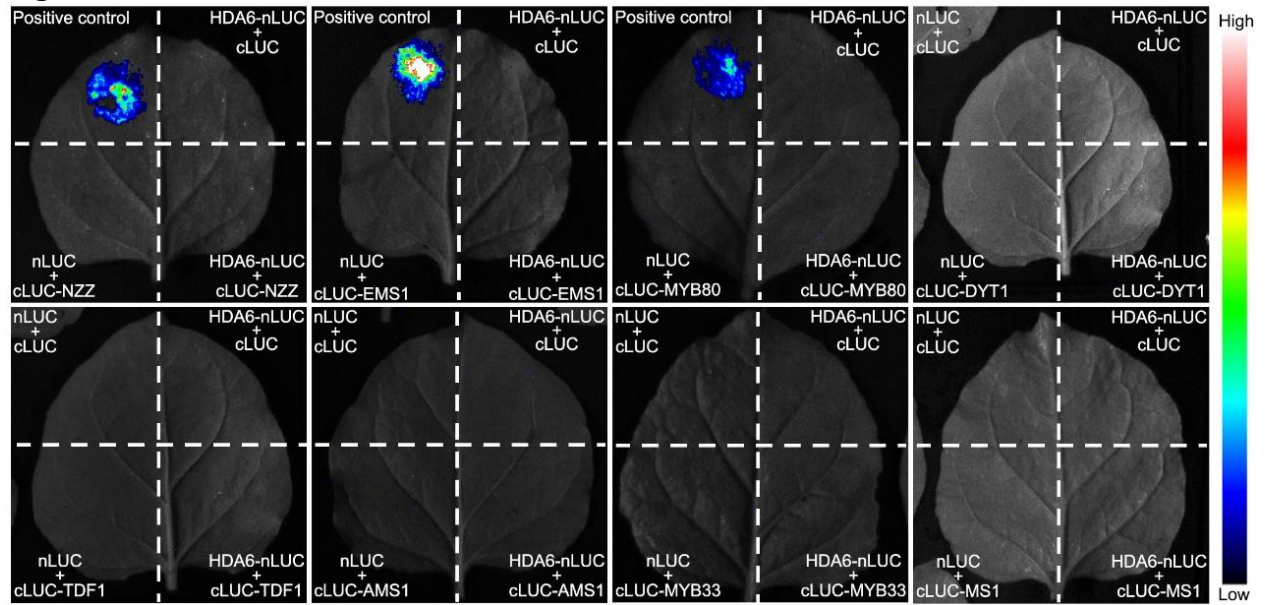

**Fig. S6. Luciferase complementation assay screening for interactions between HDA6 and candidate regulators of tapetal development and degeneration.**

Representative luciferase complementation assays in *Nicotiana benthamiana* leaves testing the interaction between HDA6-nLUC and cLUC-tagged candidate regulators involved in tapetal development or degeneration including NZZ, EMS1, MYB80, DYT1, TDF1, AMS1, MYB33 and MS1. Positive-control combinations produced strong luminescence signals, whereas no detectable luciferase complementation was observed for the tested HDA6–candidate combinations under these assay conditions. Empty-vector combinations were included as negative controls.

**Fig. S7**

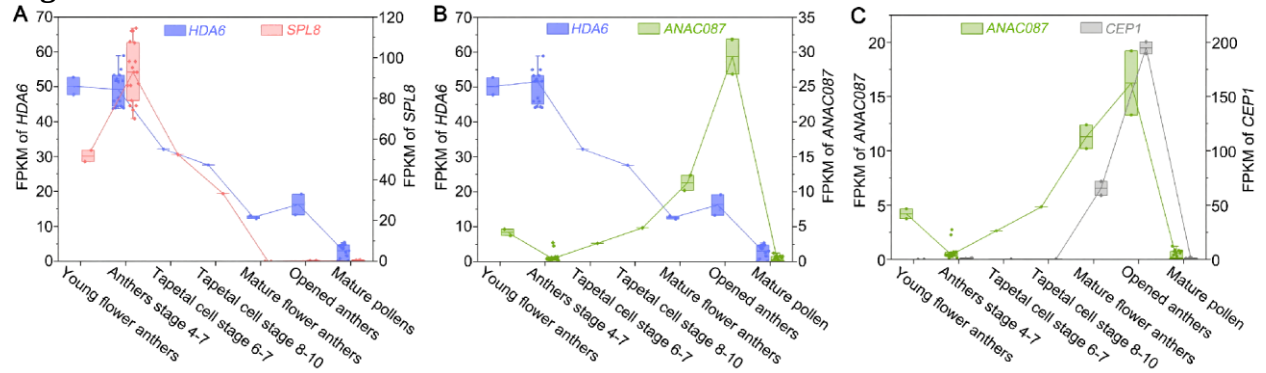

**Fig. S7. RNA-seq-based expression profile analysis of *HDA6*, *SPL8*, *ANAC087* and *CEP1* during anther development.**

(A) Transcript abundance of *HDA6* (left y-axis) and *SPL8* (right y-axis) at different anther developmental stages and in mature pollens. (B) Transcript abundance of *HDA6* (left y-axis) and *ANAC087* (right y-axis). (C) Transcript abundance of *ANAC087* (left y-axis) and *CEP1* (right y-axis). Data were obtained from publicly available datasets curated in the Arabidopsis RNA-Seq Database (ARS), including SRX1795774, SRX1796718, SRX1795824, SRX1796737, SRX1795762, SRX1796716, SRX867277, SRX867278, SRX3088664, SRX3088665, SRX275909, SRX2977304, SRX2977310, SRX2977311, GSM3402476, GSM3402477, GSM3402478, GSM3402482, GSM3402483, GSM3402484, GSM3402485, GSM3659801, GSM3659802, GSM3659803, GSM4091066, GSM4091067, GSM4091068, SRX7261876, SRX7261877, SRX638332, SRX639628, SRX639778, SRX645095, SRX645096, SRX645097, SRX645098, SRX867260, SRX867263, SRX867273, SRX867274, SRX867275, SRX867276, SRX867280, SRX867281, and SRX867283.

**Fig. S8**

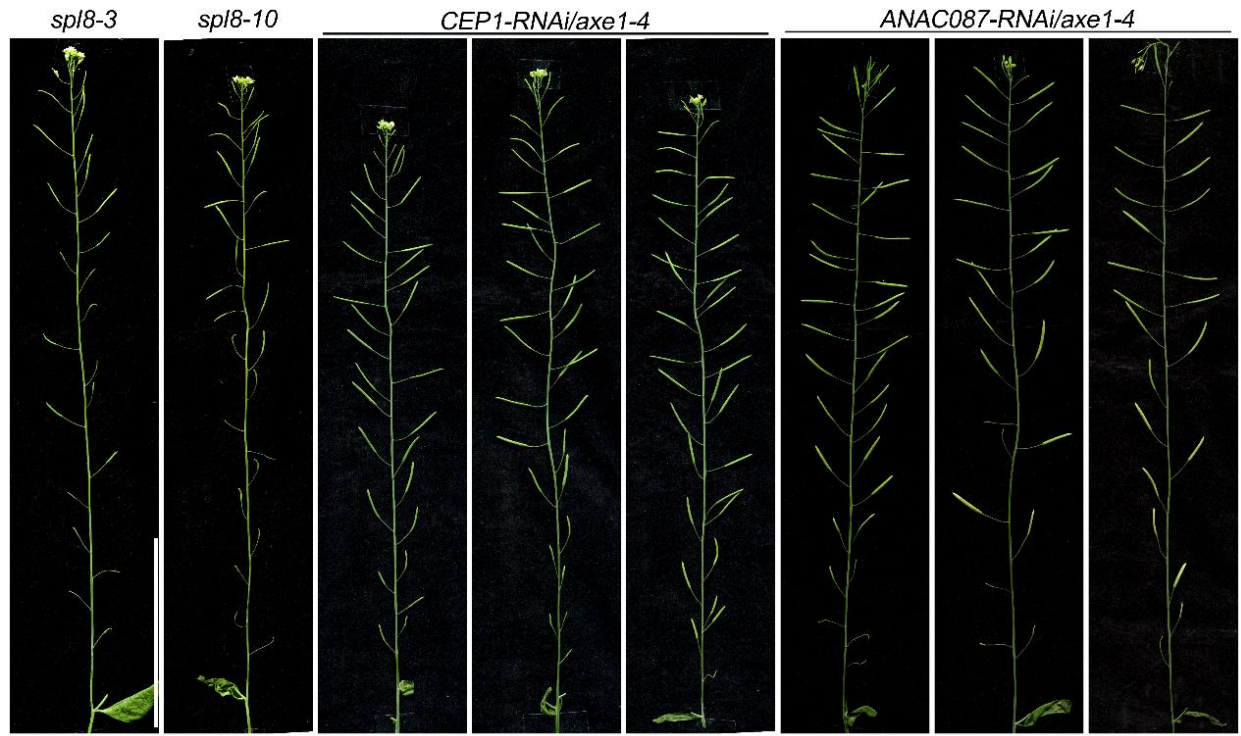

**Fig. S8. Genetic suppression of *CEP1* or *ANAC087* restores early flowering-stage fertility in *axe1-4*.**

Representative primary inflorescences of *spl8-3* and *spl8-10* mutants, and independent *CEP1-RNAi/axe1-4* and *ANAC087-RNAi/axe1-4* lines, excised below the last cauline leaf ($n \geq 3$). The *spl8* mutants exhibited severely reduced seed set during the early flowering stage, whereas RNAi-mediated suppression of *CEP1* or *ANAC087* restored silique development in the *axe1-4* background. Scale bar = 8 cm.

**Fig. S9**

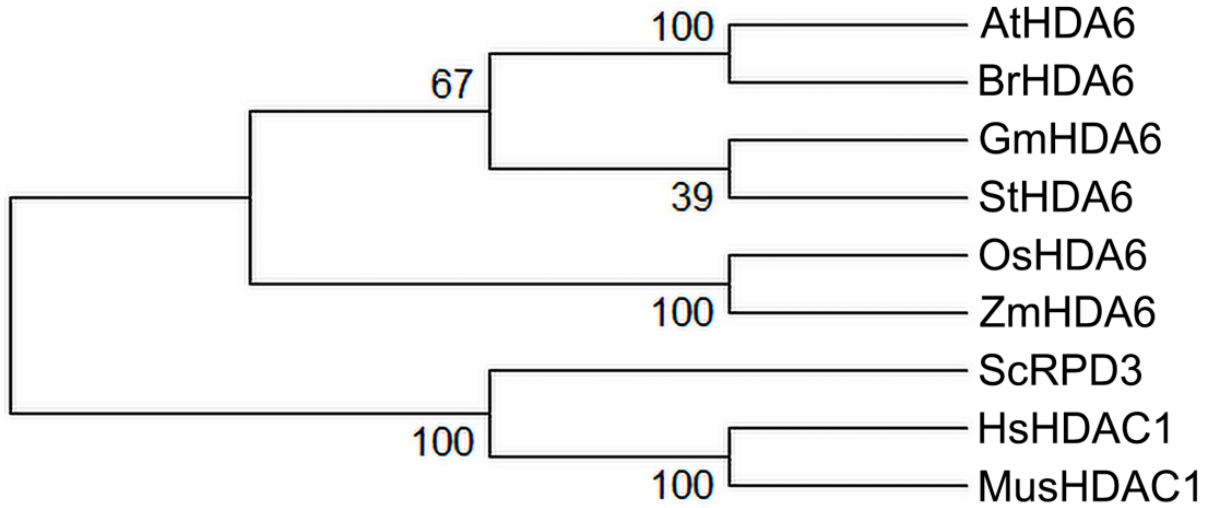

**Fig. S9. Phylogenetic analysis of HDA6 across different species.**

Maximum-likelihood phylogenetic tree generated from MUSCLE-aligned amino acid sequences of HDA6 orthologs from dicot and monocot species, together with representative fungal RPD3 and mammalian HDAC1 proteins. Arabidopsis HDA6 clusters with HDA6 orthologs from other angiosperms, forming a plant HDA6 lineage within the conserved RPD3/HDA1-type class I histone deacetylase family which is distinct from yeast RPD3 and mammalian HDAC1 proteins. Bootstrap values from 1,000 replicates are shown at the nodes. The tree was constructed in MEGA 12 using the maximum-likelihood method with the Poisson correction model. Species abbreviations: At, *Arabidopsis thaliana*; Br, *Brassica rapa*; Gm, *Glycine max*; St, *Solanum tuberosum*; Os, *Oryza sativa*; Zm, *Zea mays*; Sc, *Saccharomyces cerevisiae*; Hs, *Homo sapiens*; Mus, *Mus musculus*.
